# Impaired Proteostasis is an Early Feature of the Diabetic Heart in Humans and Mice

**DOI:** 10.64898/2026.09.01.748593

**Authors:** David R. Rawnsley, Chen Zhao, Xiucui Ma, Moydul Islam, John T. Murphy, Xumin Guan, Layla Foroughi, Attila Kovacs, Jess Nigro, Ziang Bi, Walter Navid, Honora Navid, Babak Razani, Daniel Scherr, Kartik Mani, Kenneth Margulies, L. Ashley Cowart, Simon Sedej, Ali Javaheri, Abhinav Diwan

**Author notes:** **Corresponding Author:** David Rawnsley, M.D., Ph.D.; Assistant Professor of Medicine; Washington University School of Medicine, Division of Cardiology, 660 S. Euclid Ave, CSRB-NTA Room 824, St. Louis, MO 63110. Contributed equally (Ali Javaheri, Abhinav Diwan).

## Abstract

Diabetes and obesity increase cardiac lipid levels leading to cardiomyopathy and heart failure. We hypothesized that intermittent fasting would reduce cardiac lipid levels. Surprisingly, intermittent fasting increased myocardial triglyceride content, but rescued mortality and attenuated cardiomyopathy in mice overexpressing cardiomyocyte acyl-CoA synthetase 1 (MHC-ACSL1). Lipid overload caused cardiomyocyte accumulation of polyubiquitinated protein aggregates containing desmin, a scaffolding intermediate filament protein, which intermittent fasting prevented. Furthermore, intermittent fasting reversed elevated myocardial C16:0 ceramide content, and knockdown of ceramide synthase CerS5 and CerS6 reduced palmitate-induced protein aggregation, highlighting a role for C16:0 ceramides in this pathology. Conversely, impairing aggrephagy with cardiomyocyte-specific p62 ablation induced heart failure in mice fed a high-fat diet, with paradoxically reduced cardiac lipid content. Crucially, non-failing diabetic human hearts also exhibited protein aggregate pathology. Taken together, these results demonstrate that impaired proteostasis characterizes cardiomyopathy from cardiac lipid overload and identify a promising new therapeutic target for this condition.

## INTRODUCTION

Heart failure is a leading cause of morbidity and mortality with a rising prevalence in the population ^1^. Diabetes is a significant risk factor driving this increase, as the presence of diabetes has been shown associated with a two to five-fold increase in the incidence of heart failure ^2,3^. While diabetes is traditionally associated with coronary artery disease and ischemic cardiomyopathy, epidemiologic studies have associated diabetes with increased incidence of cardiomyopathy independent of the presence of coronary artery disease ^4,5^. Among the mechanisms proposed to link diabetes and heart failure is cardiac lipotoxicity, i.e. toxic effects on cardiac cells due to the accumulation of lipid species ^6^. Further supporting this mechanistic link between cardiac lipid overload and heart failure is epidemiologic evidence that obesity in the absence of diabetes independently increases the risk of developing cardiomyopathy ^7^. Accordingly, several mouse models have been developed to induce cardiac lipid overload, using either dietary exposure to high-fat diets or transgenic modifications to promote cardiac lipid accumulation. These studies have suggested multiple potential mechanisms, such as inflammation ^8^, mitochondrial oxidative stress ^9^, and accumulation of specific toxic lipid species such as ceramides ^10,11^. Additional work is needed to further characterize these mechanisms and develop potential therapeutic strategies.

One potential intervention for lipotoxicity that has not been previously studied is intermittent fasting. Intermittent fasting involves cyclic periods of fasting followed by refeeding. Our group has previously shown that intermittent fasting improves beta cell survival and glucose tolerance in mice under high-fat diet stress ^12^. Intermittent fasting is also protective against cardiac myocyte death induced by ischemia-reperfusion injury ^13,14^ or by proteotoxic stress from a genetically mutated chaperone protein ^15^. In these studies, the benefits of intermittent fasting were shown to be mediated by stimulation of autophagy and the lysosomal biogenesis program via the transcription factor TFEB ^13,15^. As cardiomyocyte accumulation of lipids has traditionally been thought to underlie cardiac lipotoxicity, stimulation of autophagic clearance of lipids (lipophagy) is a potentially attractive therapeutic strategy. Additionally, lipotoxicity itself has been shown to impair lysosomal function and autophagy in non-cardiac tissues ^16,17^. In this scenario, stimulation of lysosomal biogenesis may help to reverse the lysosomal dysfunction induced by lipid overload,^18^ and therefore intermittent fasting may improve from cardiomyopathy induced by cardiac lipid overload through mechanisms other than a reduction in lipids.

We hypothesized that intermittent fasting would be a protective intervention in lipotoxic cardiomyopathy by facilitating clearance of accumulated lipids. To test this, we subjected MHC-ACSL1 transgenic mice, which exhibit cardiomyopathy and early mortality due to cardiac lipid accumulation ^19^, to every-other-day intermittent fasting (IF). Intermittently fasted MHC-ACSL1 mice demonstrate improved cardiac function with rescued early mortality. Examination of MHC-ACSL1 hearts revealed evidence of increased protein aggregation and impaired autophagy, and IF prevents increased protein aggregation. Dietary lipid overload also induces increased protein aggregation in the mouse heart, and IF is also protective in this context. Contrary to our hypothesis that intermittent fasting would reduce total cardiac lipid levels, IF treated animals have increased cardiac triglyceride levels versus controls, suggesting that changes in specific lipid species, rather than removal of total lipids, may underlie the benefits of IF. We subsequently show that ceramides may be the culprit lipid species. Finally, we show that diabetic non-failing human hearts also exhibit evidence of increased protein aggregation versus controls, suggesting that altered proteostasis is observed in cardiac myocytes with lipid overload in diabetics prior to development of clinical heart disease.

## METHODS

### Mice

MHC-ACSL1 mice O7 founder line ^19^ were generously provided by Dr. Jean Schaffer (Joslin Diabetes Center, Harvard University, Boston, MA). The CAG-RFP-EGFP-LC3 autophagic reporter mice ^20^ were purchased from Jackson Labs (strain number 027139). *Myh6*-Cre transgenic mice were purchased from Jackson Labs (strain number 011038). The p62 floxed allele mouse ^21^ was generously provided by Dr. Toru Yanagawa (University of Tsukuba, Japan). For all experiments, mice were backcrossed ten generations into the C57BL/6J genetic strain background. For high-fat diet mouse experiments, a 60% kcal from fat high-fat diet (Research Diets, D12492) was used in the high-fat diet arm. Control animals were fed a control low-fat chow diet with 12% kcal from fat chow diet (Lab Diet, 5053). Intermittent fasting was performed with total food deprivation and a cedar pine chip bedding change from 12:00 PM to 12:00 PM of the following day, when animals were provided access to food in alternating 24-hour cycles. Non-fasted control mice were provided ad-libitum access to fresh food. All animals were provided with ad-libitum access to water. Animals were maintained in a temperature-controlled room (22°C) on a 12 h light/dark cycle (lights on at 6:00 AM). For intermittent fasting experiments, mouse sacrifice and tissue harvesting was done between 10:00AM-12:00PM, towards the end of a 24 hour feeding period. For autophagic flux assessment, mice received an intraperitoneal injection with chloroquine (Sigma-Aldrich, CC6628; dissolved in sterile 0.9% sodium chloride, i.e. normal saline) at a dose of 80 mg/kg body weight 4 hours prior to sacrifice. Control mice underwent intraperitoneal injection with normal saline. Mice of both sexes were studied. No significant differences were observed between sexes for the primary phenotype, whereby data for both sexes were combined for presentation. Mouse studies were randomized and observers blinded. All animal studies were approved by the Institutional Animal Care and Use Committee (IACUC) at Washington University School of Medicine.

### Echocardiography

2D-directed M-mode echocardiography was performed using a Vevo 3100 Imaging System (VisualSonics) equipped with a 30 MHz linear-array transducer, as previously described ^22^. For echocardiographic studies, mice were anesthetized with 100 mg/kg intraperitoneally injected tribromoethanol (Avertin). Cardiac images were obtained by a handheld technique. Both 2-D long-axis and short axis cine loop images, as well as short-axis m-mode images, were obtained. The echocardiographer was blinded to animal genotype during image acquisition. Left ventricular dimensions, wall thickness, heart rate, and fractional shortening measurements were performed by a blinded echocardiographer using VevoStrain software (VisualSonics, Toronto, Canada).

### Histology

Histological assessment with hematoxylin & eosin (H&E) staining and Masson’s trichrome staining for fibrosis was performed as previously described ^15^. Briefly, mouse hearts were immediately dissected after euthanasia, washed in phosphate buffered saline (PBS) and then fixed overnight in 10% formalin. Fixed samples were washed with PBS, followed by dehydration with 70% ethanol. Samples were then embedded in paraffin, sectioned and mounted on glass slides, and stained with either H&E or Masson’s trichrome. Images were taken with a Zeiss Axio Imager M2 microscope.

### Electron microscopy

Transmission electron microscopy was performed on mouse hearts fixed in modified Karnovsky’s fixative, as previously described ^15^. Imaging was performed on a JEOL model 1200 EX electron microscope (JEOL, Tokyo, Japan), as previously described ^15^.

### Isolation of soluble-insoluble fractions

NP-40 insoluble fractions were isolated as described previously ^15,23^. Briefly, heart tissue was mechanically homogenized in homogenization buffer containing 0.3 M KCl, 0.1 M KH_2_PO_4_, 50 mM K_2_HPO4, 10 mM EDTA, 4 mM Na Orthovanadate, 100 mM NaF, Protease inhibitor and adjusted to pH 6.5. Homogenized samples were passed through mesh basket on ice, followed by collection of the lysate run-through which was incubated on ice for 30 minutes. A known volume of the sample was transferred to another Eppendorf tube, and 10% NP-40 was added to for a final concentration of 1% NP-40. Samples were then incubated on ice for 30 minutes, and spun at 13,000 rpm for 15 minutes, 4°C. Supernatant was collected as soluble fraction. The pellet was washed 3 times with cold PBS, with each wash followed by centrifugation at 13,000 rpm for 10 minutes. After washing, the final pellet was resuspended in 1% SDS, 10mM Tris buffer to generate the insoluble fraction, and subjected to immunoblotting as detailed below.

### Isolation of crude cardiac extracts

Crude heart extracts were isolated as previously described ^22^. Briefly, frozen heart tissue was placed in ice-cold lysis buffer (50 mM Tris HCl pH 7.4, 25 mM NaCl, 0.2% NP-40, 2.5 mM EDTA, 10 mM EGTA, 20 mM NaF, 25 mM Na_4_O_7_P_2_, 2 mM Na_3_VO_4_, supplemented with protease and phosphatase inhibitors (ThermoFisher, 78442)) and mechanically homogenized with a Qiagen TissueLyser LT homogenizer at a setting of 50 Hz for 5 minutes. Following homogenization samples were centrifuged at 300 x g for 20 minutes to remove un-lysed cells, and the supernatant was transferred to a new tube and stored at −80C. Protein concentration was determined via Bradford Assay reagent (Bio-Rad, 5000006), using a Bio-Rad SmartSpec Plus spectrophotometer.

### Immunoblotting

Immunoblotting was performed as previously described ^22,23^. Specific antibodies employed are as follows: anti-p62 (Abcam, ab56416), anti-ubiquitin (Abcam, ab134953), anti-polyubiquitinylated protein, clone FK1 (Millipore Sigma, 04-262), anti-GAPDH (Abcam, ab22555), and anti-LC3B (Novus Biologicals, NB100-2220). Protein abundance was normalized to either total protein as measured by Ponceau S staining or to GAPDH immunoblotting, as indicated in the figure.

### Immunofluorescence

Immunofluorescence studies were performed as previously described^23^. Briefly, paraffin-embedded heart sections (10 µm thick) were subjected to heat-induced epitope retrieval, followed by blocking, and incubated overnight with primary antibodies. The following primary antibodies were used: anti desmin (Santa Cruz Biotechnology, Inc, SC 7559) for mouse samples, anti-desmin (Cell Signaling, 5332) for human samples, anti p62/SQSTM1 (PROGEN Biotechnik, GP62 C), anti-ubiquitin (Abcam, ab134953), and anti-polyubiquitinylated protein, clone FK1 (Millipore Sigma, 04-262). After serial washes, samples were stained with secondary antibodies coupled to AlexaFluor488, AlexaFluor594, or AlexaFluor647 (ThermoFisher) and mounted with fluorescent 4’,6-diamidino-2-phenylindole (DAPI) mounting medium (Vector Labs, H-1200). Confocal imaging was performed on a Zeiss confocal LSM-700 laser scanning confocal microscope using 639 Zeiss Plan-Neofluar 40/1.3 and 63/1.4 oil immersion objectives, and images were acquired using Zen Blue software (Zeiss).

### Assessment of autophagic flux using the CAG-RFP-EGFP-LC3 reporter

We performed autophagic flux analyses by fluorescent microscopic examination of frozen myocardial tissue from mice with expression of the CAG-RFP-EGFP-LC3 reporters, as previously described.^22^ Briefly, mouse hearts were dissected and heart fragments were fixed for 4 hours in 4% paraformaldehyde, followed by washing with PBS. Heart tissue was then incubated overnight in 30% sucrose (dissolved in PBS). Samples were embedded in O.C.T. compound (Scigen, 4586) the following day and frozen. Tissues embedded in O.C.T. were sectioned on a Leica CM1860 UV cryostat at 10 µm thickness and were mounted on slides with VectaShield Mounting Medium with DAPI (Vector Labs, H-1200). EGFP and RFP images were acquired using a Zeiss confocal LSM-700 laser scanning confocal microscope using 639 Zeiss Plan-Neofluar 40/1.3 and 63/1.4 oil immersion objectives, and images were acquired using Zen Blue software (Zeiss). Images were then scored for RFP+GFP+ double positive puncta and for RFP+ only puncta. Only puncta associated with cells identified as cardiac myocytes based upon visualization of ‘boxcar’ shaped nuclei were counted.

### Triglyceride assay

Measurement of cardiac triglyceride levels was performed using ThermoFisher Triglyceride Reagent (Fisher Scientific, TR22421) and the following protocol. Mouse heart tissue was weighed, and ice-cold PBS was added to the tissue at a volume of 440 µl per 100 mg of heart tissue. Samples were then mechanically homogenized using a Qiagen TissueLyser LT homogenizer at a setting of 50 Hz for 5 minutes. Following homogenization, each sample was centrifuged at 600 x g and the supernatant was transferred to a new tube. The volume of each supernatant was measured and then the samples were solubilized with addition of 1% sodium deoxycholate solution at a 1:1 volume ratio, followed by incubation at 37°C for 30 minutes. Following incubation, 10 µl of solubilized sample was added to 200 µl of ThermoFisher Triglyceride Reagent in a 96-well plate and then gently mixed and was then incubated at 37°C for 15 minutes. All samples were assayed in duplicate. Following incubation, absorbance was measured at a wavelength of 540 nm using a Tecan Infinite M200 Pro plate reader. A standard curve using known triglyceride levels (Fisher Scientific, 23-666-422) was assayed at the same time as the experimental samples to convert absorbance to triglyceride concentration. All triglyceride measurements are shown normalized to input tissue weight.

### Sphingolipid profiling

Lipid extraction from frozen mouse cardiac tissue followed by sphingolipid profiling via liquid chromatography-electrospray ionization-tandem mass spectrometry (LC-ESI-MS/MS) was performed by the Virginia Commonwealth University Lipidomics and Metabolomics Core as previously described ^24–26^. Cardiac ceramide levels are shown normalized to tissue weight.

### Human tissue studies

Studies on human tissue were performed under an exemption by the IRB at Washington University School of Medicine because only de-identified human samples were used. Human heart tissue was obtained from human heart tissue banks at the University of Pennsylvania (Philadelphia, PA) and the University of Graz (Graz, Austria). All heart tissues were obtained from potential brain-dead transplant donors with no history of heart failure and were divided into two groups: individuals with and without diabetes (see Supplementary Tables S11 and S12 for clinical characteristics). After in-situ administration of cold cardioplegia via coronary perfusion, all hearts were placed on wet ice at 4°C in Krebs-Henseleit buffer.

Transmural LV samples were obtained from the LV free wall and snap frozen in liquid nitrogen with epicardial fat excluded. Isolation of soluble-insoluble fractions from human tissues was performed as described above. Immunohistochemical analyses on human tissues were performed as previously described ^23^. To quantitate the disruption of striation and increased aggregation of desmin, the following approaches were used ^23^. For striation scoring, cardiomyocytes with localization of desmin in a normal striated pattern were scored as 0, and cardiomyocytes with abnormal striation were scored as 1. For aggregate scoring, cardiomyocytes without aggregates were scored as 0 and cardiomyocytes with aggregates was scored as 1. For both striation and aggregate scoring, scores were then normalized to the total number of cardiomycoytes in the field. At least 3 images were assessed per sample. Image acquisition and quantitation were done in a blinded manner.

### Palmitate treatment in H9C2 myoblasts

Rat H9C2 cells (ATCC, CRL-1446) were cultured in DMEM supplemented with 10% fetal bovine serum, per manufacturer protocol. For siRNA experiments, On-TARGETplus SMARTPool siRNA specific to rat Cers5 (Horizon Discovery, #L-082035-02-0005) and rat Cers6 (Horizon Discovery, L-091368-02-0005) was combined at a 1:1 ratio and transfected into H9C2 cells using DharmaFect 1 Transfection Reagent (Horizon Discovery, #T-2001-01). Final siRNA concentration in culture was 25 nM. For control samples, On-TARGETplus Non-Targeting Pool (Horizon Discovery, D-001810-10-05) was used. siRNA transfection was done 48 hours prior to harvest. For palmitate experiments, cells were treated with 1 mM palmitic acid (Nu-Chek Prep, N-16-A, Elysian, MN) complexed to fatty-acid free bovine serum albumin (Lampire Biological Laboratories, 7500804) at a 2:1 ratio. Cells were treated with palmitic acid for 16 hours prior to harvest. Control samples were treated with BSA without palmitic acid. For immunoblotting experiments, cell lysates were harvested from H9C2 cells using RIPA buffer (Cell Signaling, 9806) supplemented with proteinase and phosphatase inhibitors (ThermoFisher, 78442).

### RNA isolation and quantitative reverse-transcription PCR

RNA was isolated from cultured H9C2 cells using the RNeasy Mini Kit (Qiagen, 74104). For RNA isolation from mouse cardiac tissue, the RNeasy Fibrous Tissue Mini Kit (Qiagen, 74704) was used. 1 ug of RNA was used to generate cDNA using the iScript Reverse Transcription Supermix kit (Bio-Rad, 1708841). Quantitative reverse-transcription PCR (Q-PCR) was done using iTaq Universal SYBR Green Supermix Kit (Bio-Rad, 1725121) and a QuantStudio 3 Real-Time PCR System (Thermo Fisher Scientific). All Q-PCR reactions were done in duplicate. The following primers were used for Q-PCR from mouse tissue: mCerS5 Forward 5’-CGGGGAAAGGTGTCTAAGGAT −3’, mCerS5 Reverse 5’-GTTCATGCAGTTGGCACCATT −3’, mCerS6 Forward 5’-GATTCATAGCCAAACCATGTGCC −3’, mCerS6 Reverse 5’-AATGCTCCGAACATCCCAGTC −3’, mRpl32 Forward 5’-CCTCTGGTGAAGCCCAAGATC −3’, mRpl32 Reverse 5’-TCTGGGTTTCCGCCAGTTT -3’. The following primers were used for Q-PCR from rat H9C2 cells: rCerS5 Forward 5’-ATCAGGACAAGCCTCCAACG -3’, rCerS5 Reverse 5’-AGTTGTACCAGCACTGTCGG -3’, rCerS6 Forward 5’-GGAATGAGCGGTTTTGGCTT -3’, rCerS6 Reverse 5’-ACGGCACATGGTTTGGCTAT -3’, rGAPDH Forward 5’-GGCCGAGGGCCCACT -3’, rGAPDH Reverse 5’-TGTTGAAGTCACAGGAGACAACCT -3’.

### Statistical analysis

Data are presented in graphs as Mean ± SEM. All measurements were obtained on distinct biological replicates. Statistics were performed in Prism Version 10.2.0 (GraphPad Software Inc.). Data were tested for assumptions of normality using Shapiro-Wilk Test. Statistical significance of differences was calculated via unpaired 2-tailed Student’s t-test for 2 group comparisons, and one-way ANOVA or two-way ANOVA for assessing differences between 3 or more multiple groups as indicated, followed by post-hoc testing to evaluate differences between groups. Non-parametric testing with Mann-Whitney test (for 2 groups) or Kruskal-Wallis test followed by post-hoc testing (for 3 or more groups) was utilized where indicated. Graphs containing error bars show Mean ± SEM with a p value < 0.05 considered as statistically significant. The datasets generated during and/or analyzed during the current study are available from the corresponding author on reasonable request.

## RESULTS

### Intermittent fasting reverses cardiac dysfunction and prevents early mortality in MHC-ACSL1 transgenic mice with cardiac lipid overload

To model severe cardiac lipid overload in the mouse, we studied previously described transgenic mice ^19^ with expression of acyl-CoA synthetase long chain family member 1 (ACSL1) driven by the α-myosin heavy chain (*Myh6*) promoter, henceforth referred to as MHC-ACSL1. We specifically used the O7 transgenic line which has a higher level of transgene expression, resulting in severe cardiac lipid accumulation in cardiomyocytes, cardiomyopathy, and early mortality ^19^. Based on prior work demonstrating that every-other-day intermittent fasting was cardioprotective in the setting of ischemia-reperfusion injury^13^ and protein misfolding stress,^15^ we hypothesized that intermittent fasting would also be protective in the setting of increased cardiomyocyte lipid accumulation. At 6 weeks of age, we randomized MHC-ACSL1 mice to either every-other-day intermittent fasting (IF) or an *ad-libitum* (ad-lib) diet with free access to food **(Fig. 1A)**. At this age MHC-ACSL1 mice have similar left ventricular (LV) systolic function and LV chamber size compared to wild-type mice **(Supplementary Fig. S1A-B)** but exhibit increased cardiac triglyceride levels **(Supplementary Fig. S1C)**, consistent with lipid overload. Importantly, in prior studies, we noted that mice exposed to this strategy of intermittent fasting maintain their body weight relative to control mice on an ad-lib diet,^13^ which was confirmed in MHC-ACSL1 mice **(Supplemental Table S1)**. In prior studies, we have demonstrated that intermittent fasting does not affect LV structure or function or affect myocardial histologic endpoints in wild-type mice.^13,15^ Echocardiography was performed at 12 weeks of age and survival was monitored. As previously reported, MHC-ACSL1 control mice on an ad-lib diet exhibited early mortality, with a median survival of 20.7 weeks and death of all animals by 26 weeks of age **(Fig. 1B)**. Intermittent fasting prevented early mortality in MHC-ACSL1 mice, with mice surviving up to 57 weeks of age, with a median survival of 51 weeks **(Fig. 1B)**. Intermittent fasting in MHC-ACSL1 reversed the decline in systolic function as measured by fractional shortening **(Fig. 1C, D; Supplemental Table S2)**, as well as prevented LV dilatation **(Fig. 1C, E; Supplemental Table S2)** and the increase in heart weight **(Supplemental Table S1)** and LV mass **(Fig. 1F; Supplemental Table S2)**, when compared to MHC-ACSL1 mice on an ad-lib control diet. Histological analysis showed cellular infiltration in the MHC-ACSL1 myocardium indicating myocardial inflammation as previously demonstrated in this model,^8^ and Masson’s trichrome staining showed increased myocardial fibrosis in MHC-ACSL1 mice on an ad-lib diet as compared with wild-type controls; both parameters were improved by intermittent fasting **(Fig. 1G)**. We hypothesized that the protective effect of intermittent fasting in MHC-ACSL1 mice was mediated through a reduction in lipid levels. To assess this hypothesis, we assessed cardiac triglyceride levels ^19^. Contrary to our hypothesis, intermittent fasting led to an increase in cardiac triglyceride levels relative to MHC-ACSL1 mice on an ad-libitum diet (**Fig. 1H**).

**Figure 1.**
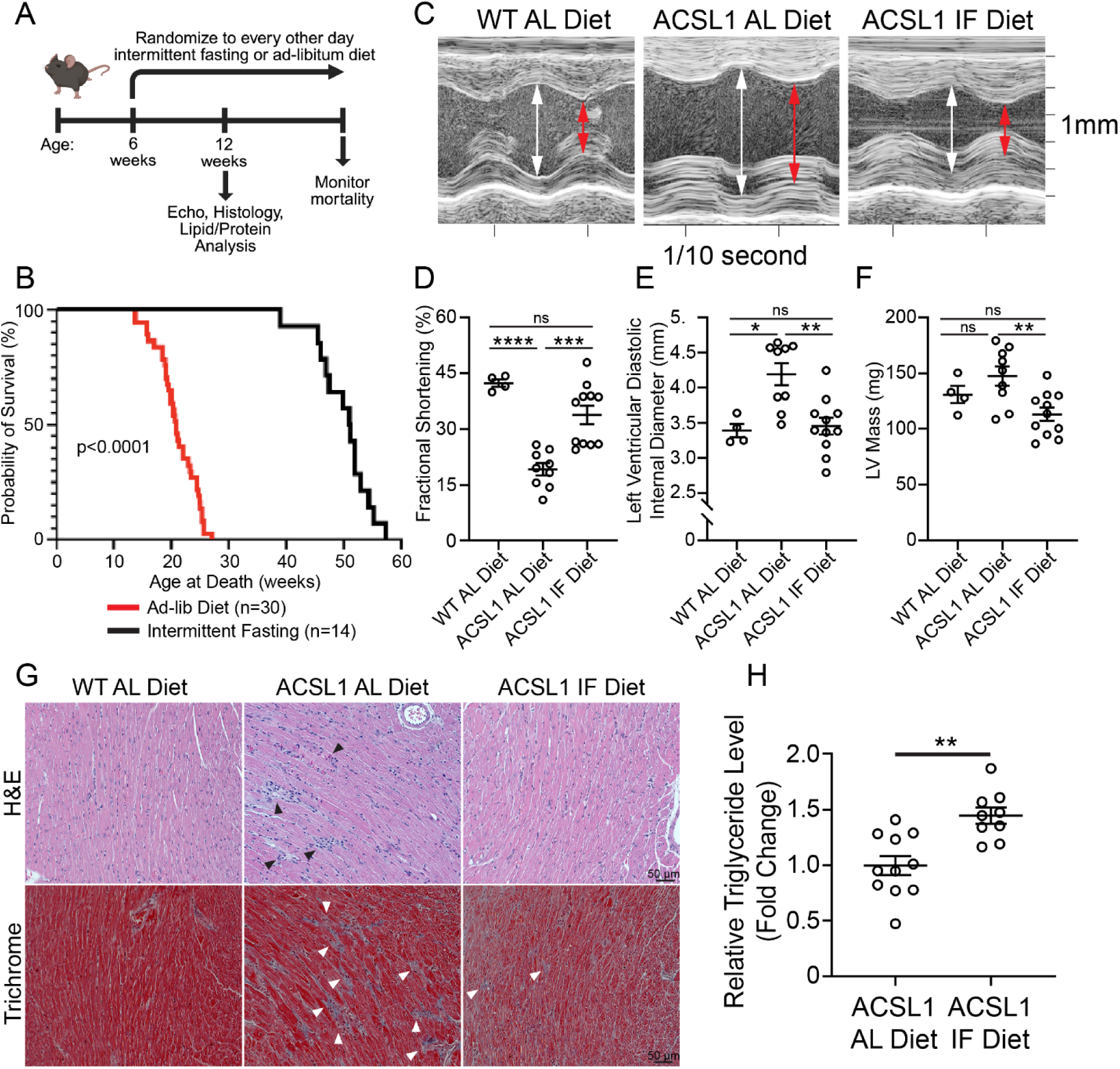
Intermittent fasting rescues cardiomyopathy and early mortality in MHC-ACSL1 mice. **(A)** Experimental strategy for intermittent fasting (IF) in MHC-ACSL1 mice. **(B)** Kaplan-Meier survival curve of MHC-ACSL1 mice on ad-lib or IF diet. P-value indicated is by log-rank test. **(C-F)** Representative 2D-directed M mode echocardiogram images (C), left ventricular fractional shortening (D), left ventricular internal diastolic diameter (E) and left ventricular mass (LVM, F) in 12-week-old wild-type (WT) and MHC-ACSL1 mice on ad-lib (AL) or intermittent fasting (IF) diets. **(G)** Hematoxylin & eosin (upper panels) and Masson’s trichrome (lower panels) staining from 12-week-old mouse hearts from indicated groups. Black arrows show areas of increased cellularity in ACSL1 hearts. White arrows show areas of fibrosis in ACSL1 hearts. **(H)** Triglyceride levels in ACSL1 hearts under AL and IF conditions. Levels shown as fold change relative to ACSL1 animals on an AL diet. Statistics by t-test (H). ns p>0.05, * p<0.05, ** p<0.01, *** p< 0.001, **** p<0.0001.

### Lipid overload induces myocardial protein aggregation in MHC-ACSL1 hearts, which is rescued by intermittent fasting

These findings indicate that the mechanism of cardiac injury in lipid-overloaded MHC-ACSL1 transgenic hearts is not mediated solely through the accumulation of triglycerides, whereby we evaluated alternative mechanisms of injury. We performed immunohistochemical staining for desmin to better characterize cardiomyocyte structure and surprisingly sections from MHC-ACSL1 mice on ad-libitum diets demonstrated disorganization of the sarcomeric structure, with mis-localization of desmin from its physiologic alignment with Z-discs and intercalated discs (see wild-type myocardium in **Fig. 2A**) to punctate accumulations that co-localized with the aggrephagy-adaptor p62 (arrowheads in MHC-ACSL1 myocardium, see **Fig. 2A**) as well as with ubiquitin (**Supplementary Fig. S2**), two markers consistent with protein aggregation ^15,23,27^.

**Figure 2.**
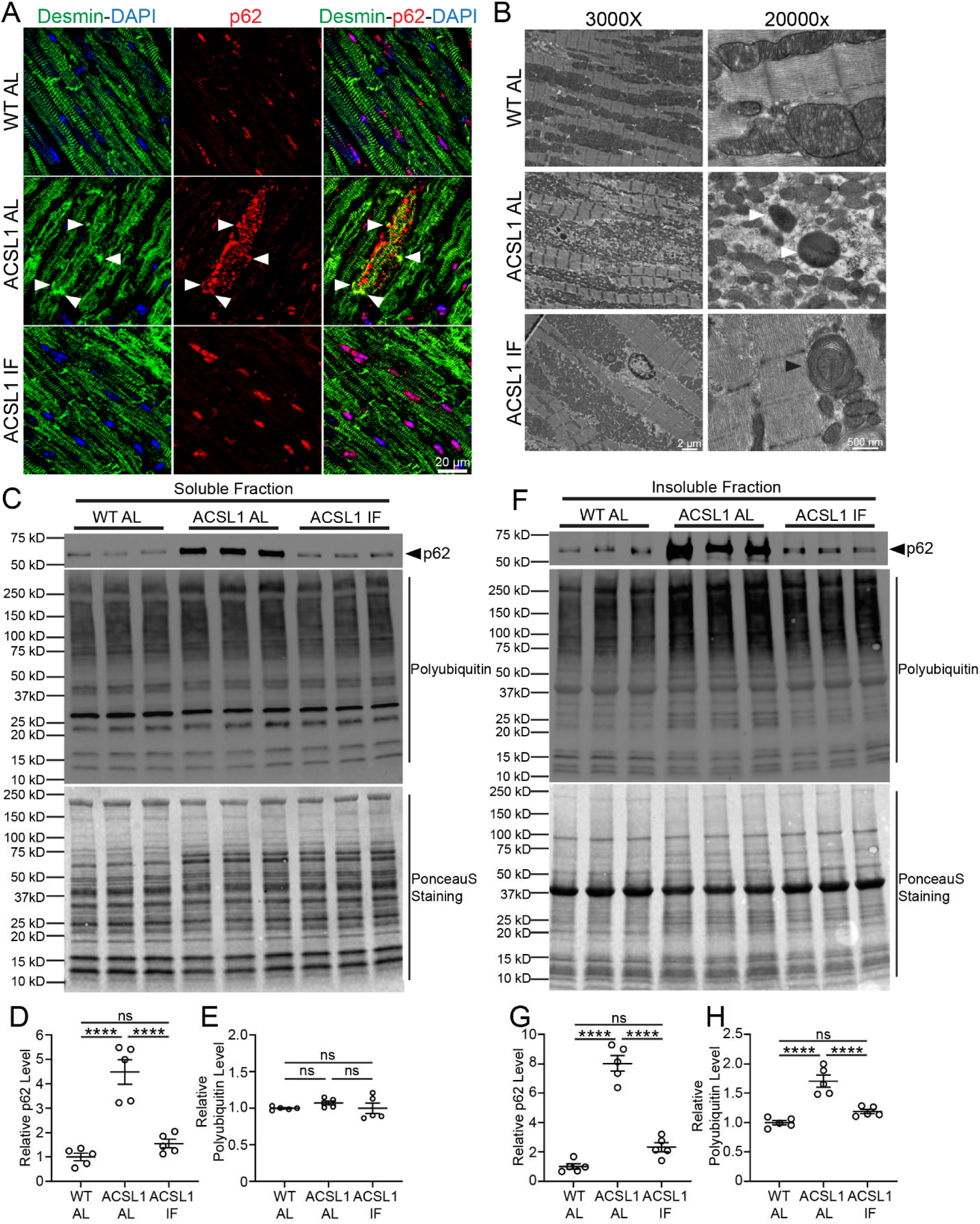
Protein aggregate pathology is observed in the myocardium of MHC-ACSL1 mice and is rescued by intermittent fasting. **(A)** Representative images demonstrating immunohistochemistry staining for desmin and p62 in myocardial sections from 12-week-old wild-type (WT) or ACSL1 transgenic hearts under ad-libitum (AL) or intermittent fasting (IF). Nuclei are stained with DAPI (blue). White arrowheads show areas of co-localization of Desmin and p62 in aggregates. **(B)** Transmission electron microscopy images of WT and ACSL1 hearts from mice treated as in A. N=2 hearts/group were examined. White arrowheads indicate electron-dense structures consistent with protein aggregates. Black arrowhead shows a mitochondrion enveloped in multiple layers of membrane. **(C-H)** Immunoblotting of the NP-40 soluble (C) and NP-40 insoluble (F) fractions from hearts of 12-week-old WT and MHC-ACSL1 mice treated as in A. Quantitation of soluble fraction p62 and polyubiquitin are shown in panel D and E, respectively. Quantitation of insoluble fraction p62 and polyubiquitin are shown in panel G and H, respectively. All samples are normalized to Ponceau S staining. Statistics in D-E and G-H are by one-way ANOVA with Tukey’s test for multiple comparison testing. ns p>0.05, **** p < 0.0001.

Desmin is an intermediate filament protein that acts as a scaffold along the Z-discs and intercalated discs to maintain normal sarcomere structure and mitochondrial localization between myofibrils,^15,28,29^ and mis-localization of desmin to the aggregates induces a state of desmin deficiency mimicking loss-of-function mutations in the *DES* gene coding for desmin.^30,31^ Desmin localization in the MHC-ACSL1 myocardium was markedly improved with intermittent fasting (**Fig. 2A, Supplementary Fig. S2**). These observations indicate that the myocardial pathology with cardiac lipid overload mimics observations with expression of R120G CRYAB mutation, and may be characterized as a desminopathy,^23,28^ that is rescued by intermittent fasting ^15^. To further characterize this phenotype, we performed transmission electron microscopy (TEM) on myocardial tissue from these mice and observed electron dense structures consistent with protein aggregates in MHC-ACSL1 hearts under ad-lib fed conditions (white arrowheads in **Fig. 2B**).

These structures were not seen in wild-type hearts and were much reduced in MHC-ACSL1 hearts under IF conditions. Electron microscopy of MHC-ACSL1 hearts showed smaller and fragmented mitochondria, and interestingly, intermittently fasted MHC-ACSL1 myocardium showed mitochondria encased in multiple layers suggesting their autophagic sequestration (**Fig. 2B)**. Taken together, these findings suggest that lipid overload in MHC-ACSL1 hearts results in disrupted proteostasis and accumulation of protein aggregates, and that intermittent fasting protects against this pathologic process.

To quantify the degree of protein aggregation in MHC-ACSL1 hearts, we homogenized heart tissue from 12-week-old MHC-ACSL1 mice under mild NP-40 detergent lysis conditions. Under these conditions, protein aggregates remain largely insoluble and are thus enriched in the insoluble fraction isolated by centrifugation ^23,32,33^. This isolated “NP-40 insoluble fraction” can then be solubilized under harsher conditions using lysis buffer containing SDS detergent, and protein aggregation abundance can be assessed by Western blotting for p62 and polyubiquitinylated conjugates. Immunoblotting of 12-week-old hearts demonstrated significantly increased levels of p62, an adaptor protein essential for protein aggregation ^34^, and polyubiquitinylated conjugates in the NP-40 insoluble fraction of ad-lib diet MHC-ACSL1 hearts versus control hearts **(Fig. 2F-H).** Intermittent fasting markedly reduced p62 and polyubiquitin levels in the insoluble fraction from MHC-ACSL1 hearts **(Fig. 2F-H).** The NP-40 soluble fraction from ACSL1 ad-lib diet mice also demonstrated an increase in p62 levels relative to controls, suggestive of increased proteotoxic stress (**Fig. 2C-E**); this phenotype was also rescued by intermittent fasting. Taken together, these findings demonstrate that increased protein aggregate accumulation occurs in response to increased cardiac lipid stress and suggest that intermittent fasting offers protection via improvement in protein quality control despite an increase in total myocardial triglyceride levels.

### Intermittent fasting reduces C16:0 ceramide levels in MHC-ACSL1 hearts

While our results demonstrate increased protein aggregation in the transgenic mouse MHC-ACSL1 heart, the mechanism through which lipids impaired protein quality control remained unclear. Given the intermittent fasting improved proteostasis in MHC-ACSL1 hearts without reducing triglyceride levels (**Fig. 1H**), we hypothesized that a specific lipid species, rather than total lipid content, mediates the toxic effects on protein quality control. Multiple lipid species have been implicated as mediators of lipotoxicity ^35^. Based on our findings of impaired proteostasis, we focused our study on ceramides, as prior work has suggested links between elevated ceramide levels and lysosomal dysfunction in both non-cardiac ^36^ and cardiac contexts ^37^. Furthermore, human studies have demonstrated associations between elevated myocardial tissue ceramide levels and worsening cardiac dysfunction,^38,39^ and inhibition of ceramide synthesis has been shown to be a cardioprotective strategy in the setting of lipid overload in mouse models^11^, although the mechanism through which ceramides exert their toxicity is unclear. To assess ceramide levels, we performed mass spectrometry on cardiac tissues from MHC-ACSL1 mice under ad-lib and intermittently fasted conditions, as well as wild-type controls. MHC-ACSL1 hearts under ad-lib dietary conditions exhibited a dramatic upregulation of C16:0 ceramide **(Fig. 3A)** versus wild-type control hearts. Intermittent fasting markedly attenuated the increase in C16:0 ceramides in MHC-ACSL1 hearts **(Fig. 3A).** Notably, there was no significant change noted in the other long chain ceramide species (C14:0, C18:1, C18:0, C20:0; **Supplementary Fig. S3A-D**). While some of very-long chain species (C24:0, C24:1, C26:0) did exhibit significant increases in MHC-ACSL1 hearts relative to WT hearts, none exhibited significant rescue with intermittent fasting (**Supplementary Fig. S3F-I**). These data suggest that increased C16:0 ceramide levels may contribute to the toxicity seen in MHC-ACSL1 hearts, and the reduction in the levels of this ceramide species may underlie the benefit observed with intermittent fasting.

**Figure 3.**
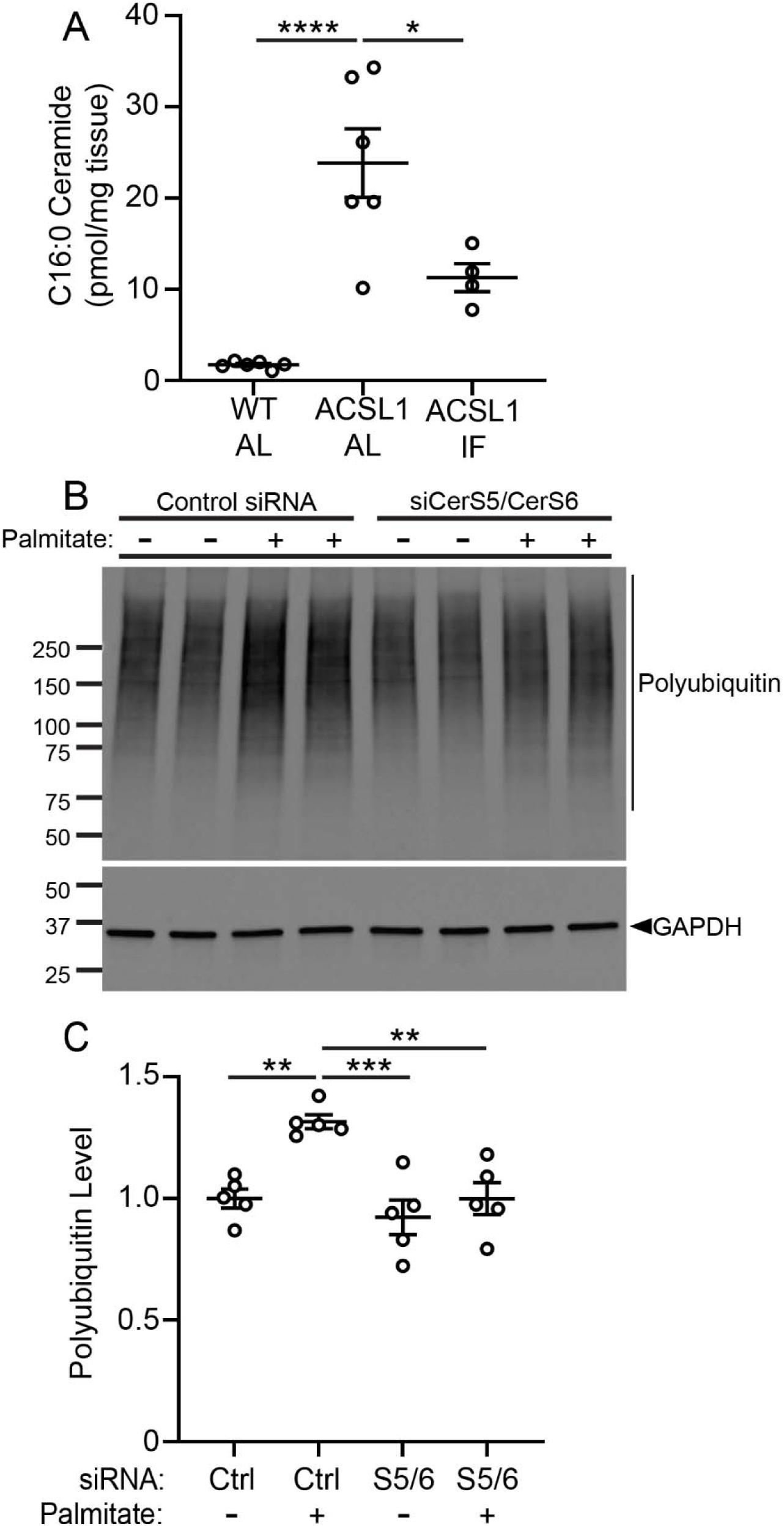
C16:0 ceramide levels are increased in MHC-ACSL1 transgenic myocardium, and knockdown of ceramide synthase 5 and ceramide synthase 6 protects against accumulation of protein aggregates in lipid overload in myocytes. **(A)** C16:0 ceramide levels from 12-week-old hearts from wild-type mice on ad-libitum diet (WT AL), MHC-ACSL1 transgenic mice on ad-libitum diet (ACSL1 AL), or MHC-ACSL1 transgenic mice after 6 weeks of every-other-day intermittent fasting (ACSL1 IF). Ceramide levels are normalized to tissue mass. Statistical comparison by one-way ANOVA followed by post-hoc testing by Tukey’s test. * p<0.05, **** p<0.0001. **(B)** Immunoblotting of H9C2 cell lysates for polyubiquitin and GAPDH following combination treatment with siRNA against ceramide synthases 5 and ceramide synthase 6 (S5/6) or control siRNA (Ctrl), and treatment with 1mM palmitate or diluent control, as indicated. siRNA treatment was initiated 48 hours prior to harvest and palmitate treatment was initiated 16 hours prior to harvest. **(C)** Quantitation of polyubiquitin levels as shown in B, normalized to GAPDH as a loading control. Statistical comparisons are by one-way ANOVA with post-hoc testing by Tukey’s test. For statistical comparisons shown, ** p<0.01, *** p<0.001, **** p<0.0001.

Therefore, we sought to directly assess whether de novo C16:0 ceramide synthesis was required for the impairment in proteostasis seen with lipid overload. Both ceramide synthase S5 (CerS5) and ceramide synthase S6 (CerS6) contribute to the synthesis of C16:0 ceramides,^40,41^ and each enzyme is upregulated in MHC-ACSL1 transgenic mouse hearts as compared with controls **(Supplementary Fig. S4A, B)**. We therefore chose to utilize a cell culture model of palmitate treatment in H9C2 rat myoblasts, which impairs autophagy.^42^ Palmitate treatment resulted in increased polyubiquitinated protein levels versus diluent control-treated cells **(Fig. 3B, C)**, consistent with impairment in proteostasis. Interestingly, siRNA treatment with knockdown of CerS5 and CerS6 **(Supplementary Fig. S4C, D)** abrogated the increase in polyubiquitinated proteins seen with palmitate treatment vs. control siRNA-treated cells **(Fig. 3B, C)**. These findings indicate that long-chain ceramide synthesis is required for the increase in polyubiquitinated aggregate-prone proteins seen with lipid overload. Taken together with in vivo assessment of ceramide levels in the MHC-ACSL1 hearts **(Fig. 3A)**, these data suggest that lipid overload impairs protein quality control through increased levels of C16:0 ceramide, and that intermittent fasting is cardioprotective through a reduction in this ceramide species.

### Intermittent fasting rescues impaired autophagy with lipid accumulation in MHC-ACSL1 cardiomyocytes

Prior work has suggested that dietary lipid overload may impair autophagy in cardiomyocytes ^43^. This led us to hypothesize that defect in protein quality control was due to impaired autophagic flux in MHC-ACSL1 mice and that intermittent fasting rescued this impairment in autophagy. To assess flux through the macro-autophagy pathway, we injected MHC-ACSL1 mice and wild type controls with chloroquine to impair lysosome acidification. Immunoblotting demonstrated accumulation of both autophagosome-bound LC3-II and the autophagic adaptor p62 only in the wild type but not in MHC-ACSL1 myocardium at 12 weeks of age, indicating impaired autophagic flux in MHC-ACSL1 mice **(Fig. 4A-C)**. Intermittently fasted MHC-ACSL1 mice demonstrated restoration of autophagic flux as compared with ad-lib fed counterparts **(Fig. 4D-F)**. As an orthogonal approach, we generated MHC-ACSL1 and control mice harboring the *CAG-RFP-EGFP-LC3* transgenic reporter allele ^20^ which drives expression of the autophagy pathway protein LC3b fused to EGFP and RFP fluorescent proteins. This tagged LC3 protein is subsequently incorporated into autophagosomes. Autophagosomes with neutral pH are labeled with both EGFP and RFP by this construct, resulting in RFP-GFP double positive LC3 puncta which are a measure of autophagosome abundance in cardiac myocytes. In acidic pH of autolysosomes, GFP fluorescence is quenched leading to RFP-only labeling, permitting assessment of autolysosome abundance. Quantitation of RFP+-GFP+ versus RFP+-only puncta therefore allows measurement of autophagosome versus autolysosome abundance, permitting assessment of autophagic flux ^20,22^. At 12 weeks of age, MHC-ACSL1 animals exhibited a decreased ratio of RFP-only/RFP-GFP puncta **(Fig. 4G-H).** This finding indicates a reduction in the number of autolysosomes relative to autophagosomes, demonstrating impairment in autophagic flux in MHC-ACSL1 hearts relative to wild-type controls. Intermittent fasting led to an increase in the ratio of RFP-only/RFP-GFP puncta in MHC-ACSL1 hearts (**Fig. 4G-H**), whereby the ratio of autophagosomes to autolysosomes was not different from wild-type myocardium. These data suggest that autophagic flux is impaired by lipid overload in MHC-ACSL1 hearts, which is rescued by intermittent fasting.

**Figure 4.**
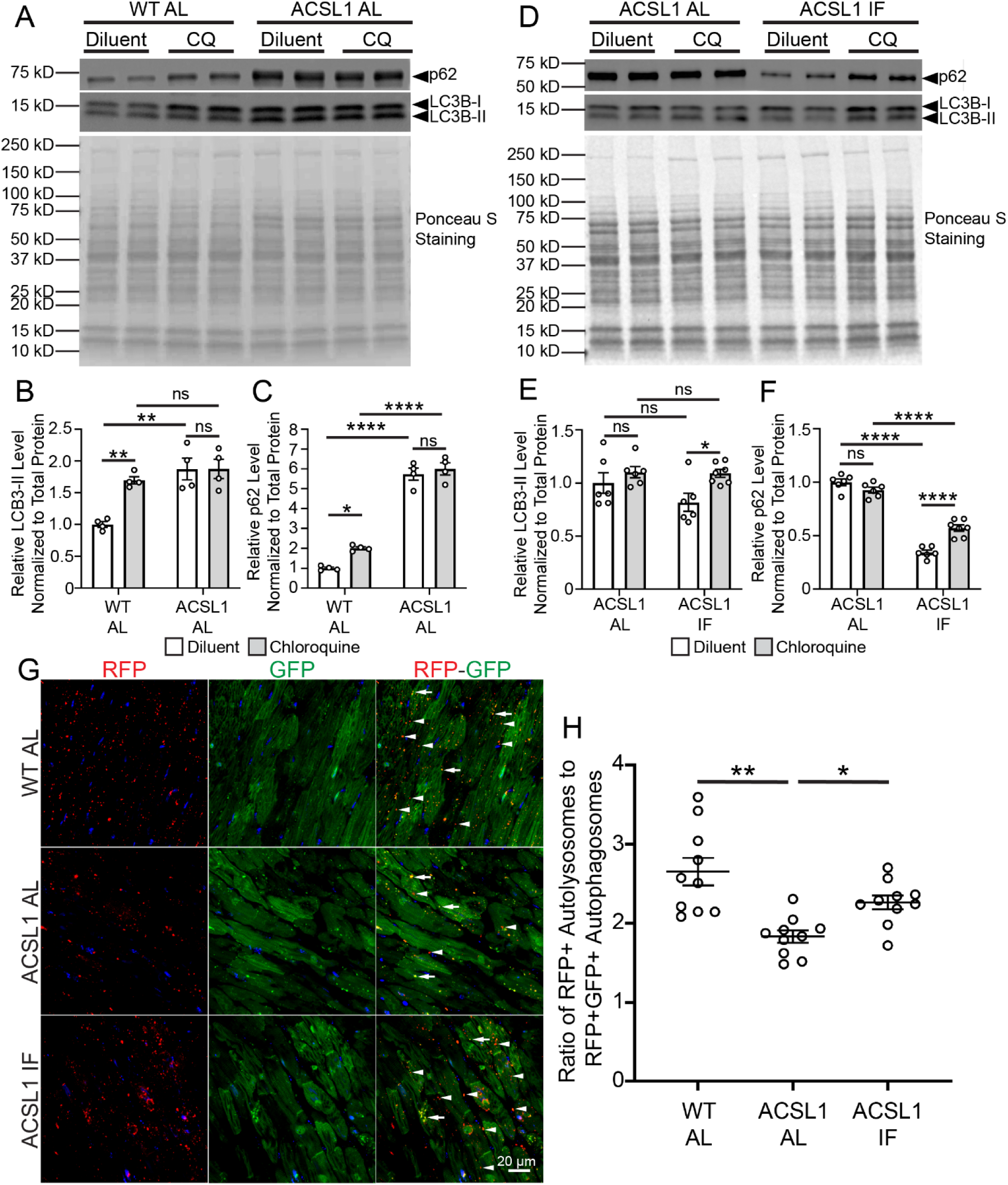
Intermittent fasting rescues impaired autophagic flux in MHC-ACSL1 transgenic hearts. **(A)** Immunoblotting for p62 and LC3B in 12-week-old wild-type (WT) and MHC-ACSL1 hearts under ad-libitum (AL) dietary conditions. Mice were treated with intraperitoneal (IP) chloroquine (CQ) injection at a dose of 80 mg/kg, or diluent injection, 4 hours prior to sacrifice. **(B-C).** Quantitation of LC3B-II (B) and p62 (C) levels as shown in A, normalized to Ponceau S staining. Statistic comparison by two-way ANOVA with Tukey’s test for multiple comparison testing. **(D)** Immunoblotting for p62 and LC3B in 12-week-old wild-type and MHC-ACSL1 hearts under ad-libitum (AL) or intermittent fasting (IF) conditions, Mice were treated with IP chloroquine (CQ) injection at a dose of 80 mg/kg, or diluent injection, 4 hours prior to sacrifice. **(E-F).** Quantitation of LC3B-II (E) and p62 (F) levels as shown in D, normalized to Ponceau S staining. Statistical comparison by two-way ANOVA with Tukey’s test for multiple comparison testing. **(G)** Fluorescent imaging of RFP and GFP expression from frozen sections from 12-week-old WT and ACSL1 mice carrying the CAG-LC3-RFP-GFP transgene under AL or IF dietary conditions for 6 weeks. White arrowheads indicate RFP+ GFP – autolysosomes; white arrows indicate double RFP+ GFP+ autophagosomes. **(H)** Ratio of RFP+ GFP-puncta (autolysosomes) to RFP+GFP+ puncta (autophagosomes) from Panel G. Puncta were scored from five images per sample and then averaged. Statistical comparison by one-way ANOVA with Tukey’s test for multiple comparison testing. For statistical comparisons in this figure, ns p>0.05, * p<0.05, ** p<0.01.

### Loss of p62, an aggrephagy adaptor, worsens lipid overload cardiomyopathy in MHC-ACSL1 mice

Our results demonstrate that the protective effects of intermittent fasting on cardiac lipotoxicity are not accompanied by reduction in triglyceride levels in MHC-ACSL1 hearts, but rather by a reduction in protein aggregation and improved autophagy. These findings suggest two possible explanations. First, these findings may indicate that cardiomyopathy seen in the MHC-ACSL1 transgenic mice may be secondary to proteotoxic stress provoked by cardiac lipid overload.

Indeed, lipid overload has been found to trigger protein misfolding in the endoplasmic reticulum to activate the unfolded protein response,^44^ and provoke protein aggregation.^45^ Alternatively, impaired proteostasis may be a secondary sequela of a dysfunctional myocardium, and not the underlying cause of cardiomyopathy. To distinguish between these two mechanisms, we chose to specifically impair aggrephagy in the MHC-ACSL1 heart by deleting the adapter p62, which is essential for protein aggregate removal by autophagy in the heart ^46,47^ and other tissues ^34,48^.

We first characterized cardiac structure and function in wild-type mice lacking p62 in cardiomyocytes. Prior publications have suggested that cardiomyocyte-specific deletion of p62 leads to reduced systolic function ^49^. We crossed the p62 conditional floxed allele to the *Myh6-* Cre line and generated cardiomyocyte-specific p62 conditional knockout mice (p62 cKO). In contrast to prior reports, p62 cKO mice did not exhibit significant change in cardiac function or structure by echocardiography at 8 weeks of age **(Supplementary Fig. S5A, B)**, and no significant histologic abnormality or fibrosis was noted by hematoxylin and eosin or Masson’s trichrome staining, respectively (**Supplementary Fig. S5C**), when compared to p62 floxed controls (p62 fl/fl). These results indicate that p62 deletion in cardiomyocytes does not cause dysfunction in young-adult hearts. We next examined macroautophagy in p62 cKO hearts. There was no impairment in autophagic flux when assessed by immuno-blotting for LC3 in p62 cKO hearts followed chloroquine injection, when compared to p62 fl/fl control mice (**Supplementary Fig. S5D-F**). We further confirmed these findings by generating p62 cKO mice bearing the CAG-RFP-EGFP-LC3 reporter allele. Assessment of autolysosome-to-autophagosome ratio showed relative increase in autolysosome abundance, suggesting increased (rather than impaired) macro-autophagy flux (**Supplementary Fig. S5G, H**). By contrast, the levels of ubiquitinated proteins were mildly increased in both the soluble and insoluble fractions in p62 ckO hearts, which (**Supplementary Fig. S5I-K)** is consistent with its role as an aggrephagy adaptor, whereby loss of p62 leads to accumulation of protein aggregates.

Together, these findings demonstrate that loss of p62 in unstressed mice does not impair systolic function or macro-autophagy, and therefore deletion of p62 is a valid strategy for further assessing the role of aggrephagy in the MHC-ACSL1 transgenic mice with cardiac lipid overload. We examined MHC-ACSL1 mice that carried *Myh6*-Cre allele and were homozygous for p62 floxed alleles (MHC-ACSL1 p62 cKO), as well as MHC-ACSL1 p62 homozygous floxed (MHC-ACSL1 p62 fl/fl) control mice, for mortality and effects on cardiac structure and function (**Fig. 5A**). MHC-ACSL1 p62 cKO mice demonstrated accelerated mortality as compared with MHC-ACSL1 p62 fl/fl control mice, with a median survival of 14 weeks versus 18 weeks (**Fig. 5B**), suggesting that loss of p62 in cardiomyocytes worsens the underlying cardiomyopathy in the MHC-ACSL1 model. We assessed cardiac function and structure by serial echocardiography at 6 weeks and 10 weeks of age. At six weeks, there was no significant difference in LV systolic function and chamber size by echocardiography between MHC-ACSL1 p62 cKO and MHC-ACSL1 p62 fl/fl mice (**Fig. 5C-D**), while LV systolic function was reduced, and left ventricle was dilated with increased heart weight and LV mass in both groups carrying the MHC-ACSL1 transgene versus wild-type controls **(Supplementary Tables S3, S4)**. However, by 10 weeks of age, there was a significant decrease in fractional shortening (**Fig. 5E**), and a significant increase in diastolic LV internal diameter (**Fig. 5F**) and increased LV mass in MHC-ACSL1 p62 cKO mice as compared with MHC-ACSL1 p62 fl/fl control mice (**Supplementary Tables S5, S6)**. These results demonstrate accelerated cardiac dysfunction in MHC-ACSL1 transgenic mice with loss of cardiomyocyte p62.

**Figure 5.**
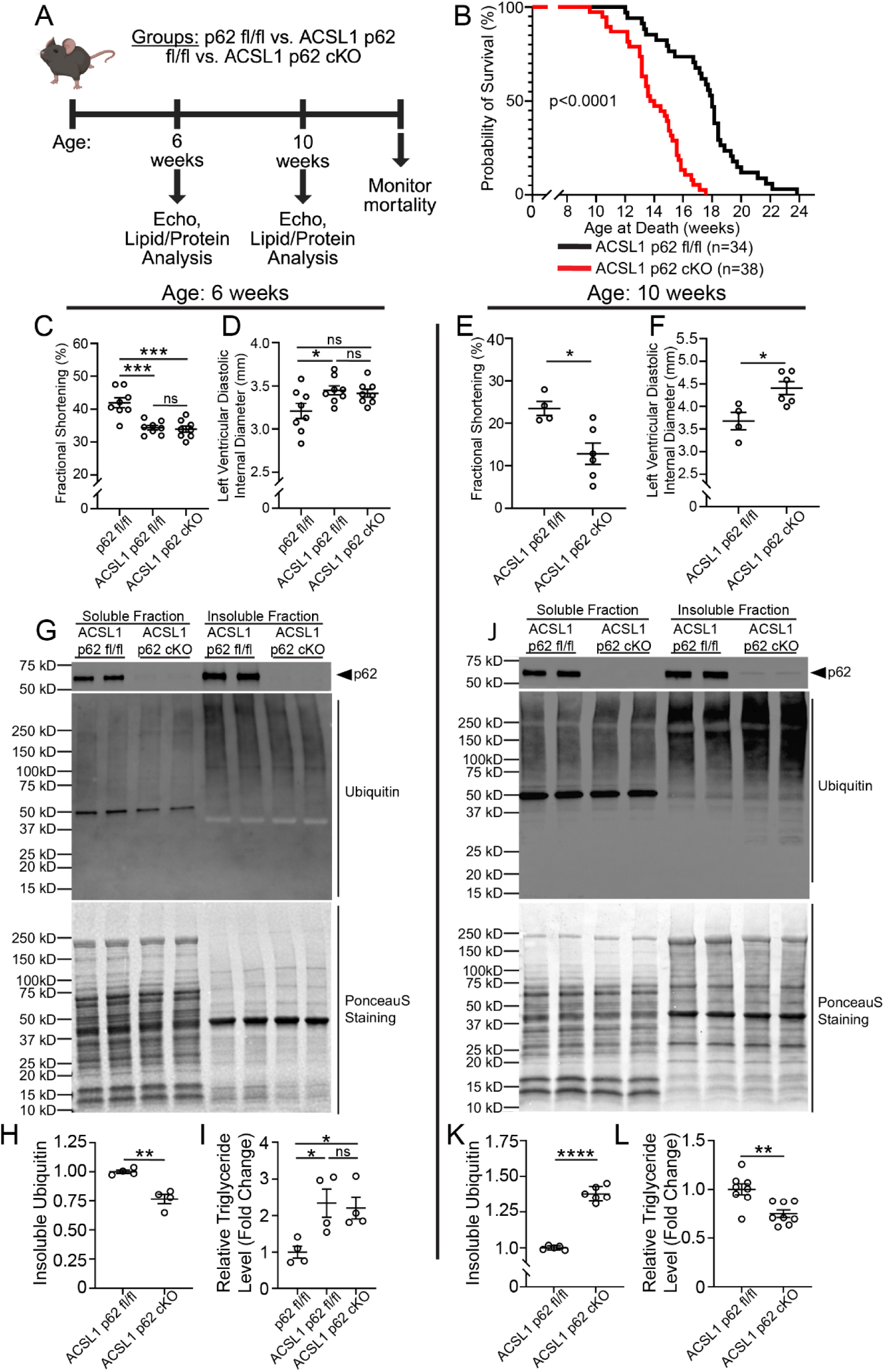
Cardiac myocyte specific ablation of p62 worsens mortality, cardiac function, and protein aggregation in MHC-ACSL1 mice without worsening lipid accumulation. **(A)** Experimental plan to assess mortality and cardiac function in p62 flox/flox (p62 fl/fl) mice, MHC-ACSL1 p62 flox/flox (ACSL1 p62 fl/fl) mice, and MHC-ACSL1-*Myh6*-Cre p62 flox/flox (ACSL1 p62 cKO) mice. **(B)** Survival of MHC-ACSL1 p62 fl/fl vs. ACSL1 p62 cKO mice. P-value shown is by log-rank test. **(C-D)** Left ventricular endocardial fractional shortening (C) and left ventricular diastolic internal diameter (D) by echocardiography in p62 fl/fl vs. MHC-ACSL1 p62 fl/fl vs. MHC-ACSL1 p62 cKO mice at 6 weeks of age. Statistical testing by one-way ANOVA with Tukey’s test for multiple comparison testing. **(E-F)** Left ventricular endocardial fractional shortening (E) and left ventricular diastolic internal diameter (F) by echocardiography in MHC-ACSL1 p62 fl/fl vs. MHC-ACSL1 p62 cKO mice at 10 weeks of age. Statistical testing by t-test. **(G)** Immunoblotting of the NP-40 soluble and NP-40 insoluble fractions from 6-week-old MHC-ACSL1 p62 fl/fl vs. MHC-ACSL1 p62 cKO mice. **(H)** Quantitation of ubiquitin levels in the insoluble fraction from panel G. Ubiquitin levels are normalized to total protein as measured by Ponceau S staining. Statistical testing by t-test. **(I)** Triglyceride levels in 6-week-old hearts from indicated genotypes. Values are normalized to mean value of p62 fl/fl control hearts. Statistical testing by one-way ANOVA with Tukey’s test for multiple comparison testing. **(J)** Immunoblotting of the NP-40 soluble and NP-40 insoluble fractions from 10-week-old MHC-ACSL1 p62 fl/fl vs. MHC-ACSL1 p62 cKO mice. **(K)** Quantitation of ubiquitin levels in the insoluble fraction from panel J. Ubiquitin levels are normalized to total protein as measured by Ponceau S staining. Statistical testing by t-test. **(L)** Triglyceride levels in 10-week-old hearts from indicated genotypes. Values are normalized to mean value of MHC-ACSL1 p62 fl/fl hearts. Statistical testing by t-test. For statistical comparisons shown in this figure, ns p>0.05, * p <0.05, ** P <0.01, *** p<0.001, **** p<0.0001.

To determine the underlying cause of cardiac dysfunction with loss of p62, we examined the levels of protein aggregation at each of these time-points. Prior to the onset of cardiac dysfunction at 6 weeks of age, there was no increase in ubiquitinated proteins in the insoluble fraction of hearts from MHC-ACSL1 p62 cKO mice relative to MHC-ACSL1 p62 fl/fl control mice (**Fig. 5G, H**). In contrast, with the onset of significant cardiac dysfunction by 10 weeks of age, ubiquitinated proteins in the insoluble fraction are increased in MHC-ACSL1 p62 cKO hearts relative to MHC-ACSL1 p62 fl/fl control myocardium (**Fig. 5J, K**). Importantly, triglyceride levels are similar between MHC-ACSL1 p62 cKO and MHC-ACSL1 p62 fl/fl mice at 6 weeks of age (**Fig. 5I**) and reduced in MHC-ACSL1 p62 cKO mice relative to controls at 10 weeks of age (**Fig. 5L**) indicating that the progressively worsening cardiomyopathy with p62 ablation was not due to increased cardiac lipids.

To test the hypothesis that the accelerated cardiomyopathy and increased protein aggregation is due to worsening macro-autophagy in MHC-ACSL1 p62 cKO hearts, we utilized the CAG-RFP-EGFP-LC3 reporter allele to assess autophagic flux at 10 weeks of age. As previously observed **(see Fig. 4A-C)**, MHC-ACSL1 p62 fl/fl mice exhibited impaired autophagic flux relative to p62 fl/fl control mice (**Supplementary Fig. S6A, B**). Notably, MHC-ACSL1 p62 cKO mice did not exhibit worsening of autophagic flux relative to MHC-ACSL1 p62 fl/fl controls (**Supplementary Fig. S6A, B**). Taken together, these results demonstrate that deletion of protein aggrephagy adaptor protein p62 in cardiomyocytes results in accelerated cardiomyopathy and earlier mortality in ACSL1 mice and implicates worsening aggrephagy as the underlying mechanism.

To examine whether p62 plays a mechanistic role in the beneficial effects of intermittent fasting on cardiac lipotoxicity, we subjected MHC-ACSL1 p62 fl/fl and MHC-ACSL1 p62 cKO mice to intermittent fasting (IF) or ad-libitum (AL) diet at 6 weeks of age (**Fig. 6A**). Strikingly, left ventricular fractional shortening and chamber size were markedly improved with IF in MHC-ACSL1 p62 cKO mice at 10 weeks of age (**Fig. 6B, C; Supplementary Table S7**), indicating that IF was protective against the accelerated cardiomyopathy seen in these animals. Reflecting this improvement in cardiac function, MHC-ACSL1 p62 cKO mice under intermittent fasting conditions also demonstrated improvement in mortality (**Fig. 6D**), with an increase in median survival from 14 to 33 weeks when compared to ad-lib fed (AL) controls. While there was clear benefit with improved mortality upon intermittent fasting in MHC-ACSL1 p62 cKO mice, their median survival was shorter as compared with the median survival of intermittently fasted MHC-ACSL1 control animals (51 weeks in ACSL1 IF, see **Fig. 1B**). This was accompanied by reduced left ventricular fractional shortening and left ventricular dilation in intermittently fasted MHC-ACSL1 p62 cKO versus intermittently fasted MHC-ACSL1 p62 fl/fl mice at 28 weeks of age (**Fig. 6E, F; Supplementary Table S8**). These findings show that intermittent fasting prolongs lifespan in MHC-ACSL1 transgenic mice via both p62-dependent aggrephagy and p62-independent mechanisms.

**Figure 6.**
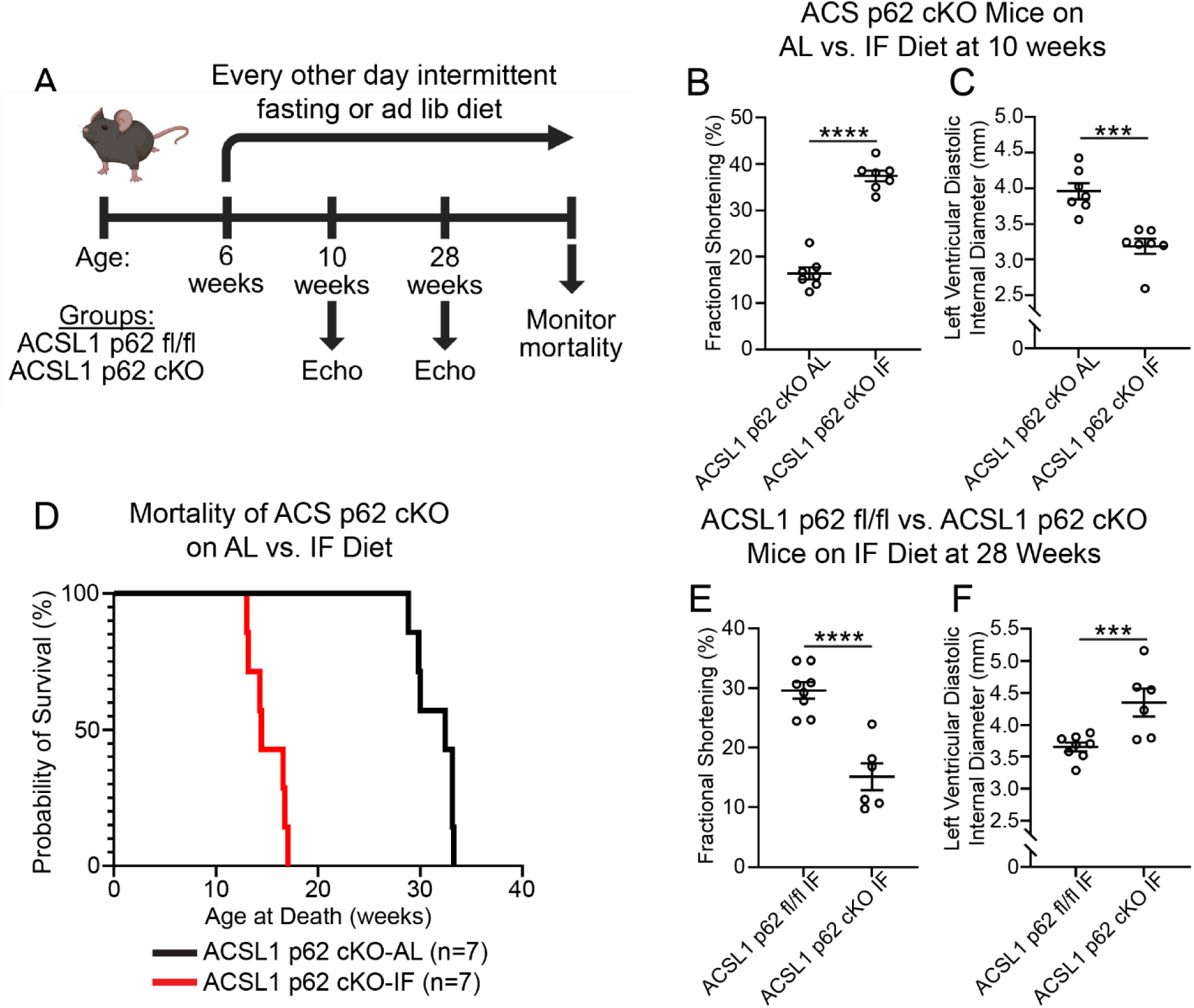
Cardiac myocyte specific ablation of p62 abrogates the beneficial effects of intermittent fasting in ACSL1 mice. **(A)** Experimental plan to assess the effect of intermittent fasting on MHC-ACSL1 p62 cKO and MHC-ACSL1 p62 fl/fl mice, randomized to every-other-day intermittent fasting (IF) or ad-libitum (AL) diets at 6 weeks of age, with assessment of cardiac function by echocardiography at 10 weeks and 28 weeks of age. **(B-C)** Left ventricular endocardial fractional shortening (B) and left ventricular diastolic internal diameter (C) by echocardiography in MHC-ACSL1 p62 cKO mice at 10 weeks of age. Statistical testing by t-test. **(D)** Survival of MHC-ACSL1 p62 cKO mice under the indicated dietary interventions. P-value shown is for log rank testing. **(E-F)** Left ventricular endocardial fractional shortening (E) and left ventricular diastolic internal diameter (F) by echocardiography in intermittently fasted (IF) MHC-ACSL1 p62 fl/fl and MHC-ACSL1 p62 cKO mice at 28 weeks of age. Statistical testing by t-test. For statistical comparisons shown in this figure, *** p<0.001, **** p<0.0001.

### Loss of p62 provokes cardiomyopathic decompensation in mice with diet-induced obesity

To determine if dietary lipid overload also induces myocardial protein aggregate pathology, we fed mice with a high-fat diet (HFD). Previous studies have demonstrated that HFD administration results in cardiac hypertrophy and diastolic dysfunction, particularly when combined with L-NAME therapy.^50^ However, C57BL/6J mice do not manifest systolic dysfunction with HFD feeding alone.^51^ We hypothesized that HFD treatment in WT mice induces protein aggregation in the heart (as observed in the MHC-ACSL1 myocardium) and that impaired aggrephagy with p62 ablation will result in cardiomyopathy with systolic dysfunction. To test this hypothesis, we provided ad-lib access to HFD (60% kcal from fat) or a control chow diet to p62 cKO mice and p62 fl/fl control mice for 24 weeks starting at 8 weeks of age, followed by assessment of endpoints at 32 weeks of age (**Fig. 7A**). HFD treatment in p62 fl/fl mice induced weight gain and increased heart weight (normalized to tibial length) **(Supplementary Table S9)** without affecting LV systolic function **(Fig. 7B; Supplementary Table S10)** or LV chamber size (**Fig. 7C, D; Supplementary Table S10**) versus chow fed controls. Mice with cardiomyocyte deletion of p62 on a chow diet (p62 cKO chow) did not demonstrate differences in body weight, heart weight or cardiac structure and function as compared with p62 fl/fl controls at 32 weeks of age (**Fig. 7B-D**). In contrast, p62 cKO animals treated with HFD also gained body weight with increased heart weight as compared with chow fed p62 cKO mice (**Supplementary Table S9**) but manifested a significant reduction in LV systolic function (**Fig. 7B; Supplementary Table S10**) relative to p62 cKO on a control chow diet and to p62 fl/fl control mice fed HFD. Moreover, p62 cKO mice on HFD also developed a significant increase in left ventricular systolic chamber diameter (**Fig. 7D**), a trend towards increased LV ventricular diastolic chamber diameter (**Fig 7C**) and increased left ventricular mass as compared with HFD-fed p62 fl/fl mice despite equivalent increase in body weight and heart weight (**Supplementary Table S9)**, consistent with adverse cardiac remodeling induced by cardiac myocyte p62 ablation under lipid overload stress. Histological assessment of myocardial sections from these mice revealed increased fibrosis in p62 cKO mice on HFD, relative to either p62 fl/fl mice on HFD or p62 cKO mice on control chow (**Supplementary Fig. S7A**). Examination of myocardial ultrastructure with TEM imaging of p62 cKO mice on HFD demonstrated marked accumulation of protein aggregates (**Supplementary Fig. S7B**).

**Figure 7.**
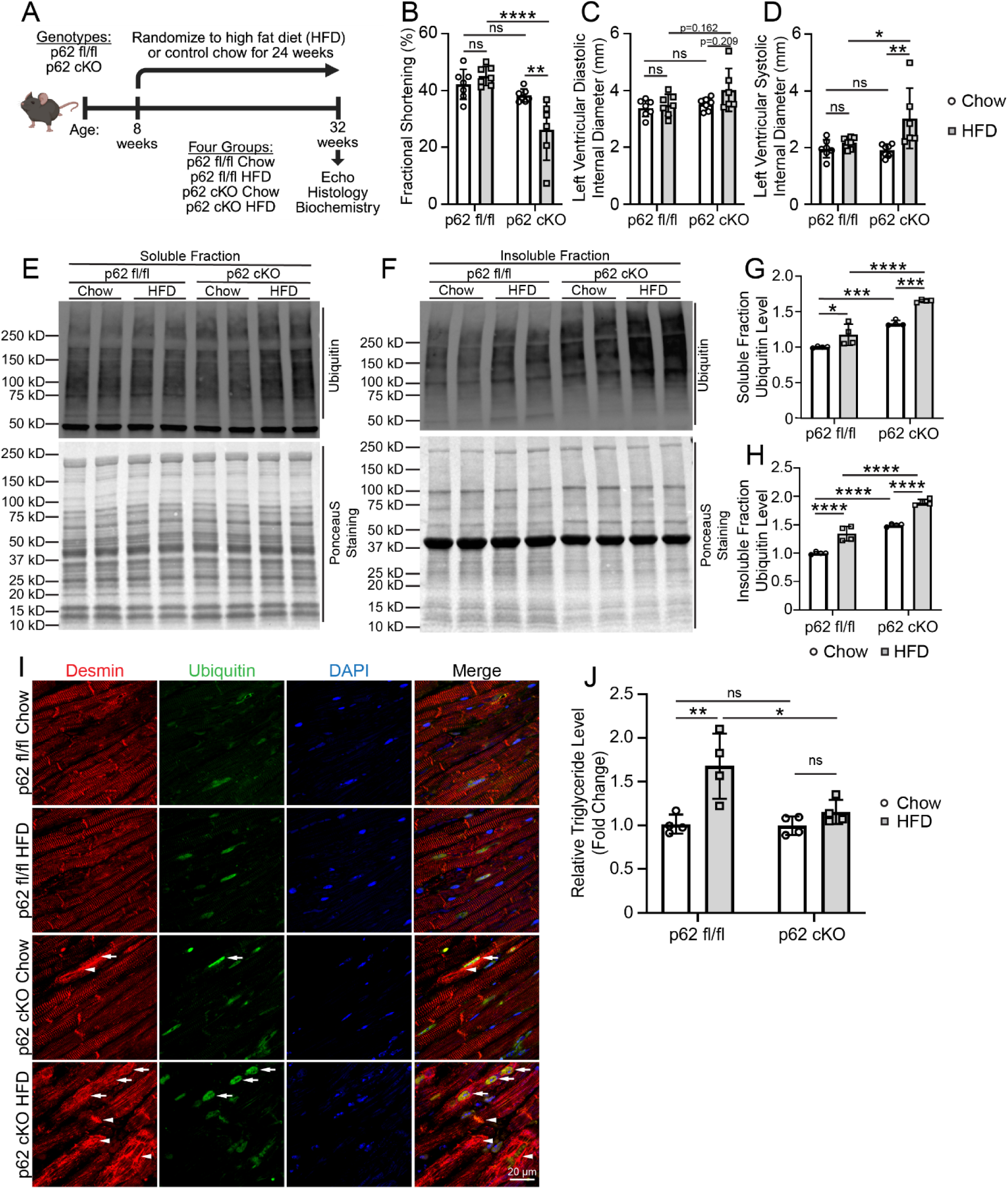
High fat diet feeding induces myocardial protein aggregate pathology and loss of p62 induces cardiomyopathic decompensation with high fat diet. **(A)** Experimental plan to assess the effect of high fat diet feeding on p62 homozygous floxed (fl/fl) and *Myh6*-Cre p62 flox/flox (p62 cKO) as compared with chow fed controls. **(B-D)** Left ventricular endocardial fractional shortening (B), left ventricular diastolic internal diameter (C), and left ventricular systolic internal diameter (D) in p62 cKO and p62 fl/fl mice fed a high fat diet or control chow for 24 weeks as in A. Statistical testing by two-way ANOVA with Tukey’s test for multiple comparison testing. **(E-H)** Representative immunoblots showing the NP-40 soluble (E) and NP-40 insoluble fractions (F) from myocardium of 32-week-old p62 cKO and p62 fl/fl mice fed a high fat diet or control chow for 24 weeks as in A. Quantitation of ubiquitin levels in the soluble (G) and insoluble fractions (H) from panels E and F respectively. Ubiquitin levels are normalized to total protein as measured by Ponceau S staining. Statistical testing by two-way ANOVA with Tukey’s test for multiple comparison testing. **(I)** Immunohistochemistry staining of 32-week-old myocardial sections from p62 cKO and p62 fl/fl mice fed a high fat diet or control chow for 24 weeks as in A. Images show staining for Desmin (red), ubiquitin (green), and DAPI (blue). Arrow points to protein aggregates. **(J)** Triglyceride levels in hearts from 32-week-old p62 cKO and p62 fl/fl mice fed a high fat diet or control chow for 24 weeks as in A. Levels shown as fold change relative to p62 fl/fl mice on a chow diet. Statistical testing by two-way ANOVA with Tukey’s test for multiple comparison testing. *p<0.05, **p<0.01, *** p<0.001, **** p<0.0001.

We next performed soluble-insoluble biochemical fractionation on heart samples followed by immunoblotting. In both the soluble and insoluble fractions, HFD increased the levels of ubiquitinated proteins in p62 fl/fl control mice, and this was further increased in p62 cKO mice on HFD treatment (**Fig. 7E-H**), consistent with impaired proteostasis in these hearts. Notably, p62 fl/fl mice fed a HFD diet did not exhibit sarcomeric abnormalities or obvious protein aggregates by desmin and ubiquitin staining (**Fig. 7I**), despite an increase in polyubiquitinated proteins (**Fig. 7E-H**). In contrast, p62 ablation in cardiac myocytes resulted in accumulation of desmin in aggregates with relative preservation of sarcomeric structure, which was worsened with mis-localization of desmin from its physiologic location on Z discs and intercalated discs to desmin-containing aggregates that co-localized with polyubiquitinated proteins in HFD-fed p62 cKO myocardium (see arrows in **Fig. 7I**). Remarkably, while p62 fl/fl mice exhibited the expected increase in cardiac triglyceride levels under HFD conditions relative to chow control (**Fig. 7J**), p62 cKO mice fed a HFD diet did not exhibit a significant increase in triglyceride levels **(Fig. 7J**) despite weight gain and increase in heart weight (**Supplementary Table S9)** as compared to chow p62 cKO mice. These findings mimic the reduced total triglycerides seen in MHC-ACSL1 p62 cKO mice versus MHC-ACSL1 p62 fl/fl controls (**Fig. 5L**) demonstrating that the systolic dysfunction observed in models of cardiac lipid overload is closely related to accumulation of protein aggregates rather than lipid accumulation.

### Diabetic human hearts from individuals without known heart disease exhibit increased protein aggregation

Our findings demonstrate that impaired proteostasis and protein aggregate accumulation characterize both a genetic and a dietary mouse model of cardiac lipid overload. To extend these inquiries to humans, we examined heart tissue from diabetic patients for evidence of increased protein aggregation. We excluded patients with known heart failure to avoid confounding, and both our diabetic and non-diabetic control samples were taken from patients with no known clinical heart failure. We performed soluble-insoluble biochemical fractionation of these myocardial tissue samples, followed by immunoblotting for ubiquitin and p62. There were no significant changes seen in the soluble samples (data not shown). However, p62 levels were increased in the insoluble fraction from diabetic samples relative to controls (**Fig. 8A, B**), as were the levels of polyubiquitinated proteins (**Fig. 8A, C**). Immunohistochemical analyses revealed disorganization of desmin (**Fig. 8D, E**) with protein aggregates where desmin co-localized with polyubiquitinated proteins (**Fig. 8D, F**) only in the diabetic myocardium. These findings demonstrate that myocardial lipid stress in humans results in accumulation of protein aggregates in human diabetic myocardium, even in the absence of heart failure.

**Figure 8.**
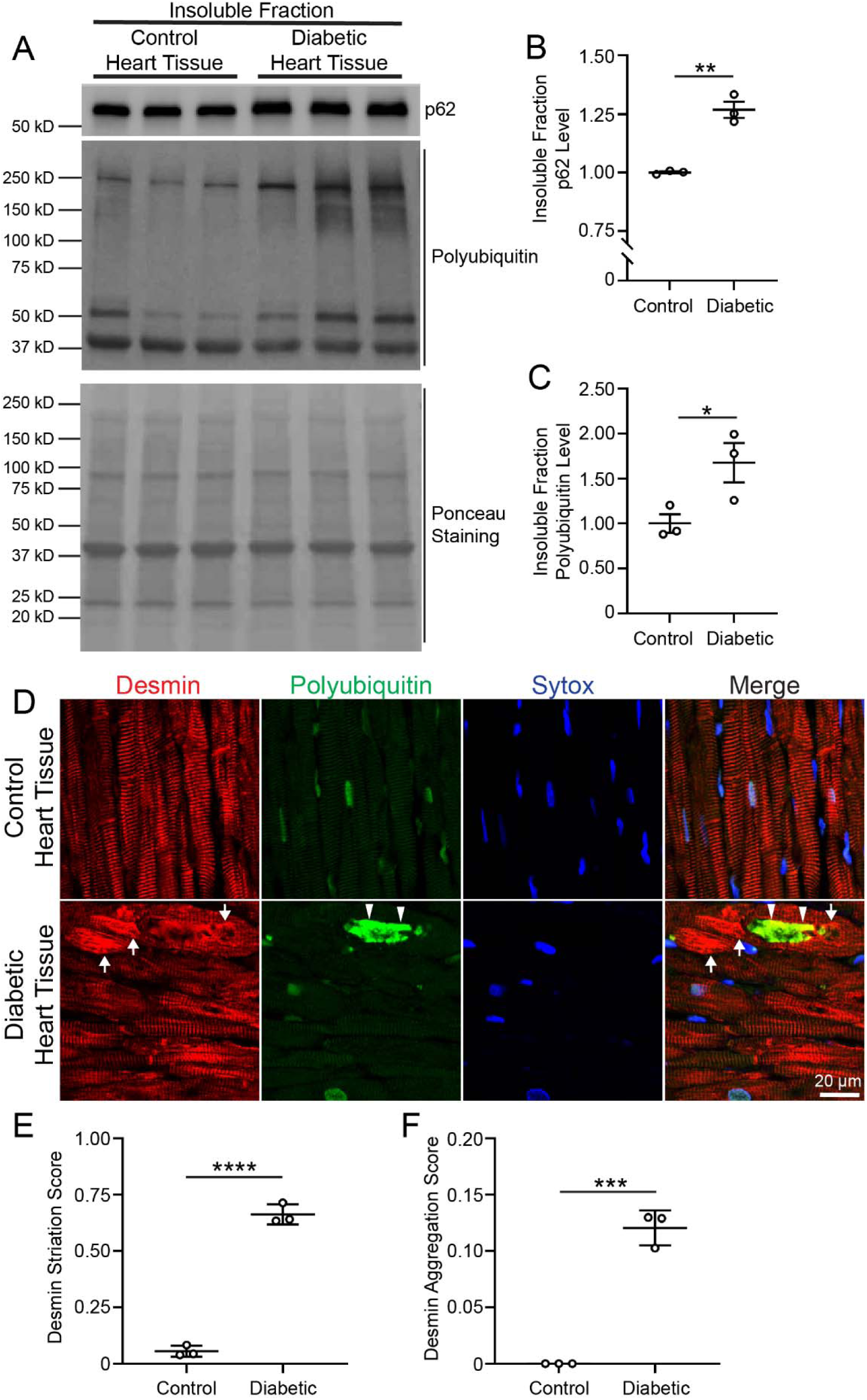
Diabetes induces protein aggregate pathology in non-failing human hearts. **(A)** Western blot for polyubiquitinylated proteins and p62 in the NP-40 insoluble fraction from heart tissue from human diabetic and non-diabetic (control) patients. **(B-C)** Level of p62 (B) and polyubiquitinylated proteins (C) in A, normalized to total protein as measured by Ponceau S staining. Statistics by unpaired t-test. ** p<0.01, *** p<0.001. **(D)** Immunohistochemical staining for Desmin (red), polyubiquitin (green), and Sytox stain to identify nuclei (blue). Arrows indicate examples of desmin aggregation, arrowheads indicated polyubiquitin aggregation. **(E-F)** Quantitation of the percentage of cardiac myocytes with disorganized Desmin striation pattern (E) and Desmin aggregates (F) from images as done in panel D. See methods for description of quantitative methods. Five images were taken per sample and then averaged. N=3 samples per group. Statistics by unpaired t-test. *** p<0.001, **** p<0.0001.

## DISCUSSION

Our study demonstrates that intermittent fasting is a protective strategy in lipotoxic cardiomyopathy and prevents early mortality in transgenic MHC-ACSL1 mice. We also find that the cardiac lipid overload in these transgenic mice is characterized by increased protein aggregation, and that intermittent fasting is protective against protein aggregation. Surprisingly, intermittent fasting does not reduce total lipid levels, but rather alters the levels of ceramides, particularly ceramide C16:0, and cognate ceramide synthases appear necessary to drive protein aggregation in myocytes. Remarkably, impairment of aggrephagy is sufficient to provoke left ventricular systolic dysfunction in mice fed with myocardial lipid overload triggered by high fat diet. Finally, we show that increased protein aggregation is also observed in human diabetic hearts under increased lipid stress without heart failure. Taken together, these data suggest that altered protein quality control is a pathogenic mechanism for cardiomyopathy under lipid overload stress.

Our study yields several novel findings. First, cardiac protein aggregate pathology is increased in lipid overload in both transgenic MHC-ACSL1 mice and in dietary lipid overload and this pathology can be ameliorated by intermittent fasting. Furthermore, human hearts exhibit similar pathology under lipid stress from diabetes, suggesting this extends to human patients as well. Prior evidence linking impaired cardiac proteostasis to cardiomyopathy has generally been limited to rare inherited cardiomyopathies due to genetic mutations in chaperone proteins such as CRYAB ^15,27^ and BAG3 ^52^. These findings potentially extend the relevance of altered cardiac proteostasis as a pathophysiologic mechanism to cardiomyopathy secondary to obesity and diabetes, which is much more common. Our data indicate that autophagy is altered in MHC-ACSL1 mice under increased lipid stress and thus decreased autophagic clearance of aggregates leads to accumulation. However, we cannot rule out the possibility of additional disruptions of other mechanisms of aggregate removal (such as the proteasome), as well as the possibility that protein misfolding is increased by lipotoxic stress and contributes to increased protein aggregation. Further studies will be needed to dissect these possible mechanisms.

Contrary to our hypothesis, our measurements of triglyceride levels in MHC-ACSL1 hearts under ad-lib and IF conditions definitively show that intermittent fasting does not reduce cardiac triglyceride accumulation. Instead, our data suggest that alterations in certain lipid species, specifically ceramides, underlie both the toxicity of lipid overload and the protective effects of intermittent fasting in these models. These findings are supported by prior studies which demonstrated that overexpression of diacylglycerol acyltransferase 1 (DGAT1), which catalyzes triacylglycerol formation from diacylglycerol and fatty acyl-CoA, improved cardiac dysfunction in a mouse model of cardiac lipotoxicity, while increasing cardiac triglyceride levels and reducing ceramide levels ^53^. It is intriguing that intermittent fasting may have a similar mechanism of action in the heart, potentially favoring the storage of accumulated lipids in relatively benign triglyceride forms rather than toxic species such as ceramides. Ceramide 16:0 in particular was significantly increased in MHC-ACSL1 ad-lib hearts versus controls, with a reduction in levels upon intermittent fasting. This finding is interesting in light of human epidemiological studies, which have linked elevated plasma levels of ceramide 16:0 to increased risk of developing heart failure ^54,55^. Further studies are needed to definitively link ceramide 16:0 to cardiomyopathy in our mouse models. As individual ceramide synthase isoforms have different N-acyl chain length specificities ^56^, it would potentially be possible to reduce cardiac ceramide C16:0 levels by targeting specific ceramide synthases which are involved in the synthesis of long-chain ceramides like C16:0 ^10,56^.

The finding that intermittent fasting can rescue cardiomyopathy and early mortality in MHC-ACSL1 mice has important translational implications. Intermittent fasting has previously been studied in clinical trials in human diabetic patients and has been shown to be an effective strategy for improving obesity, insulin resistance, and metabolic parameters such as low-density lipoprotein and total cholesterol levels ^57,58^. However, studies have not yet examined whether intermittent fasting can improve clinical cardiovascular outcomes in these populations, and further clinical studies are needed to address this hypothesis. Furthermore, the benefits of intermittent fasting related to stimulation of lysosomal biogenesis and autophagy can potentially be replicated with pharmacological approaches, which may be easier to implement in clinical populations than a dietary lifestyle intervention like intermittent fasting. As a candidate, the non-reducing disaccharide trehalose has been shown to activate lysosomal biogenesis in a beneficial manner in multiple cardiovascular conditions ^59–61^.

A remarkable finding of our studies is that in failing hearts, total myocardial triglyceride levels are directly proportional to left ventricular systolic function (see Figure panels 1D and H; 5E and L; and 7B and J); i.e. cardiac lipid levels increase with better systolic function and vice versa. This raises the intriguing possibility that while lipid overload may trigger myocardial dysfunction earlier in the disease process, impaired protein quality control is the driving force for progression of myocardial dysfunction, and the failing myocardium may increasingly rely on triglyceride stores as a source of energy. Indeed, restoring long chain fatty acyl CoA-driven mitochondrial metabolism mitigates cardiac lipotoxicity by restoring the balance of cardioprotective C20- and C22-ceramides and reducing cardiotoxic C16-ceramides ^62^. Overall, our dataset points to a role for the quality of lipids, i.e. specific lipid species versus the quantity (i.e. total abundance) of lipids as a driver of myocardial pathology and points to a critical downstream mechanistic role for protein aggregation therein. One such putative mechanism is the generation of aggregate-prone proteins via toxic effects of ceramides, which then induce protein aggregation and sequester normal proteins such as desmin in the aggregates, provoking proteotoxicity downstream of lipid overload. This is analogous to the mechanism that we have recently demonstrated where CRYAB, a cardiac chaperone protein becomes aggregate prone by persistent p38 kinase driven phosphorylation in post-myocardial infarction myocardium and sequesters normal sarcomeric and sarcomere-associated proteins in aggregates including cardiac myocyte dysfunction^23,33^. Analogously, targeting the proteotoxicity by preventing formation of aggregate-prone proteins, as we achieved with 25-hydroxycholesterol treatment^23^ or accelerating the removal of aggregate-prone proteins and protein aggregates by stimulating the autophagy-lysosome machinery^15^ are strategies that need to be explored in countering cardiac lipotoxicity in future studies.

## Acknowledgements

This study was supported by R01 grants from the National Institutes of Health (HL107594 and HL143431 and) and the Department of Veterans Affairs (I01BX004235, 1I01BX005981) to A.D. D.R.R. was supported by a T32HL007081 and K08HL163469 from the National Institutes of Health and X.G. was supported by a post-doctoral research award from the American Heart Association. We are grateful to Wandy Beatty, Ph.D., Department of Molecular Biology at Washington University School of Medicine for assistance with electron microscopy studies. We also wish to thank Joan Avery at the Center for Cardiovascular Research at Washington University School of Medicine, for her administrative assistance with conducting the study.

## Disclosure statement

A.D. reports consulting for clinical trials with Clario (previously ERT/Biomedical systems); and A.D. and K.M. served on the scientific advisory board for Dewpoint Therapeutics, which did not affect the current study. Other authors do not have any competing interests to report.

## Notes

### Competing Interest Statement

Abhinav Diwan reports consulting for clinical trials with Clario (previously ERT/Biomedical systems); and Abhinav Diwan and Kartik Mani served on the scientific advisory board for Dewpoint Therapeutics, which did not affect the current study. Other authors do not have any competing interests to report.

## References

1. Palaniappan, L.P., Allen, N.B., Almarzooq, Z.I., Anderson, C.A.M., Arora, P., Avery, C.L., Baker-Smith, C.M., Bansal, N., Currie, M.E., Earlie, R.S., et al. (2026). 2026 Heart Disease and Stroke Statistics: A Report of US and Global Data From the American Heart Association. Circulation 153, e275–e906. 10.1161/cir.0000000000001412.

2. Kannel, W.B., Hjortland, M., and Castelli, W.P. (1974). Role of diabetes in congestive heart failure: the Framingham study. Am J Cardiol 34, 29–34.

3. Nichols, G.A., Hillier, T.A., Erbey, J.R., and Brown, J.B. (2001). Congestive heart failure in type 2 diabetes: prevalence, incidence, and risk factors. Diabetes Care 24, 1614–1619. 10.2337/diacare.24.9.1614.

4. Iribarren, C., Karter, A.J., Go, A.S., Ferrara, A., Liu, J.Y., Sidney, S., and Selby, J.V. (2001). Glycemic control and heart failure among adult patients with diabetes. Circulation 103, 2668–2673. 10.1161/01.cir.103.22.2668.

5. Pazin-Filho, A., Kottgen, A., Bertoni, A.G., Russell, S.D., Selvin, E., Rosamond, W.D., and Coresh, J. (2008). HbA 1c as a risk factor for heart failure in persons with diabetes: the Atherosclerosis Risk in Communities (ARIC) study. Diabetologia 51, 2197–2204. 10.1007/s00125-008-1164-z.

6. Ritchie, R.H., and Abel, E.D. (2020). Basic Mechanisms of Diabetic Heart Disease. Circ Res 126, 1501–1525. 10.1161/CIRCRESAHA.120.315913.

7. Kenchaiah, S., Evans, J.C., Levy, D., Wilson, P.W., Benjamin, E.J., Larson, M.G., Kannel, W.B., and Vasan, R.S. (2002). Obesity and the risk of heart failure. N Engl J Med 347, 305–313. 10.1056/NEJMoa020245.

8. Schilling, J.D., Machkovech, H.M., Kim, A.H., Schwendener, R., and Schaffer, J.E. (2012). Macrophages modulate cardiac function in lipotoxic cardiomyopathy. American journal of physiology. Heart and circulatory physiology 303, H1366–1373. 10.1152/ajpheart.00111.2012.

9. Tsushima, K., Bugger, H., Wende, A.R., Soto, J., Jenson, G.A., Tor, A.R., McGlauflin, R., Kenny, H.C., Zhang, Y., Souvenir, R., et al. (2018). Mitochondrial Reactive Oxygen Species in Lipotoxic Hearts Induce Post-Translational Modifications of AKAP121, DRP1, and OPA1 That Promote Mitochondrial Fission. Circ Res 122, 58–73. 10.1161/circresaha.117.311307.

10. Russo, S.B., Baicu, C.F., Van Laer, A., Geng, T., Kasiganesan, H., Zile, M.R., and Cowart, L.A. (2012). Ceramide synthase 5 mediates lipid-induced autophagy and hypertrophy in cardiomyocytes. J Clin Invest 122, 3919–3930. 10.1172/JCI63888.

11. Park, T.S., Hu, Y., Noh, H.L., Drosatos, K., Okajima, K., Buchanan, J., Tuinei, J., Homma, S., Jiang, X.C., Abel, E.D., and Goldberg, I.J. (2008). Ceramide is a cardiotoxin in lipotoxic cardiomyopathy. J Lipid Res 49, 2101–2112. 10.1194/jlr.M800147-JLR200.

12. Liu, H., Javaheri, A., Godar, R.J., Murphy, J., Ma, X., Rohatgi, N., Mahadevan, J., Hyrc, K., Saftig, P., Marshall, C., et al. (2017). Intermittent fasting preserves beta-cell mass in obesity-induced diabetes via the autophagy-lysosome pathway. Autophagy 13, 1952–1968. 10.1080/15548627.2017.1368596.

13. Godar, R.J., Ma, X., Liu, H., Murphy, J.T., Weinheimer, C.J., Kovacs, A., Crosby, S.D., Saftig, P., and Diwan, A. (2015). Repetitive stimulation of autophagy-lysosome machinery by intermittent fasting preconditions the myocardium to ischemia-reperfusion injury. Autophagy 11, 1537–1560. 10.1080/15548627.2015.1063768.

14. Ahmet, I., Wan, R., Mattson, M.P., Lakatta, E.G., and Talan, M. (2005). Cardioprotection by intermittent fasting in rats. Circulation 112, 3115–3121. 10.1161/CIRCULATIONAHA.105.563817.

15. Ma, X., Mani, K., Liu, H., Kovacs, A., Murphy, J.T., Foroughi, L., French, B.A., Weinheimer, C.J., Kraja, A., Benjamin, I.J., et al. (2019). Transcription Factor EB Activation Rescues Advanced alphaB-Crystallin Mutation-Induced Cardiomyopathy by Normalizing Desmin Localization. Journal of the American Heart Association 8, e010866. 10.1161/JAHA.118.010866.

16. Liu, H.Y., Han, J., Cao, S.Y., Hong, T., Zhuo, D., Shi, J., Liu, Z., and Cao, W. (2009). Hepatic autophagy is suppressed in the presence of insulin resistance and hyperinsulinemia: inhibition of FoxO1-dependent expression of key autophagy genes by insulin. J Biol Chem 284, 31484–31492. 10.1074/jbc.M109.033936.

17. Yang, L., Li, P., Fu, S., Calay, E.S., and Hotamisligil, G.S. (2010). Defective hepatic autophagy in obesity promotes ER stress and causes insulin resistance. Cell Metab 11, 467–478. 10.1016/j.cmet.2010.04.005.

18. Rawnsley, D.R., and Diwan, A. (2020). Lysosome impairment as a trigger for inflammation in obesity: The proof is in the fat. EBioMedicine 56, 102824. 10.1016/j.ebiom.2020.102824.

19. Chiu, H.C., Kovacs, A., Ford, D.A., Hsu, F.F., Garcia, R., Herrero, P., Saffitz, J.E., and Schaffer, J.E. (2001). A novel mouse model of lipotoxic cardiomyopathy. The Journal of clinical investigation 107, 813–822. 10.1172/JCI10947.

20. Li, L., Wang, Z.V., Hill, J.A., and Lin, F. (2014). New autophagy reporter mice reveal dynamics of proximal tubular autophagy. J Am Soc Nephrol 25, 305–315. 10.1681/ASN.2013040374.

21. Harada, H., Warabi, E., Matsuki, T., Yanagawa, T., Okada, K., Uwayama, J., Ikeda, A., Nakaso, K., Kirii, K., Noguchi, N., et al. (2013). Deficiency of p62/Sequestosome 1 causes hyperphagia due to leptin resistance in the brain. The Journal of neuroscience : the official journal of the Society for Neuroscience 33, 14767–14777. 10.1523/jneurosci.2954-12.2013.

22. Ma, X., Rawnsley, D.R., Kovacs, A., Islam, M., Murphy, J.T., Zhao, C., Kumari, M., Foroughi, L., Liu, H., Qi, K., et al. (2022). TRAF2, an Innate Immune Sensor, Reciprocally Regulates Mitophagy and Inflammation to Maintain Cardiac Myocyte Homeostasis. JACC Basic Transl Sci 7, 223–243. 10.1016/j.jacbts.2021.12.002.

23. Islam, M., Rawnsley, D.R., Ma, X., Navid, W., Zhao, C., Guan, X., Foroughi, L., Murphy, J.T., Navid, H., Weinheimer, C.J., et al. (2025). Phosphorylation of CRYAB induces a condensatopathy to worsen post-myocardial infarction left ventricular remodeling. The Journal of clinical investigation 135. 10.1172/jci163730.

24. Shaner, R.L., Allegood, J.C., Park, H., Wang, E., Kelly, S., Haynes, C.A., Sullards, M.C., and Merrill, A.H., Jr. (2009). Quantitative analysis of sphingolipids for lipidomics using triple quadrupole and quadrupole linear ion trap mass spectrometers. J Lipid Res 50, 1692–1707. 10.1194/jlr.D800051-JLR200.

25. Haynes, C.A., Allegood, J.C., Park, H., and Sullards, M.C. (2009). Sphingolipidomics: methods for the comprehensive analysis of sphingolipids. J Chromatogr B Analyt Technol Biomed Life Sci 877, 2696–2708. 10.1016/j.jchromb.2008.12.057.

26. Kovilakath, A., Mauro, A.G., Valentine, Y.A., Raucci, F.J., Jamil, M., Carter, C., Thompson, J., Chen, Q., Beutner, G., Yue, Y., et al. (2024). SPTLC3 Is Essential for Complex I Activity and Contributes to Ischemic Cardiomyopathy. Circulation 150, 622–641. 10.1161/CIRCULATIONAHA.123.066879.

27. Rawnsley, D.R., Islam, M., Zhao, C., Guan, X., Kargar Gaz Kooh, Y., Mendoza, A., Navid, H., Kumari, M., Pandi, P., Murphy, J.T., et al. (2026). Mitophagy Facilitates Cytosolic Proteostasis to Preserve Cardiac Function. Circ Res. 10.1161/circresaha.126.328328.

28. Goldfarb, L.G., and Dalakas, M.C. (2009). Tragedy in a heartbeat: malfunctioning desmin causes skeletal and cardiac muscle disease. J.Clin.Invest 119, 1806–1813. 38027 [pii];10.1172/JCI38027 [doi].

29. Tsikitis, M., Galata, Z., Mavroidis, M., Psarras, S., and Capetanaki, Y. (2018). Intermediate filaments in cardiomyopathy. Biophys Rev 10, 1007–1031. 10.1007/s12551-018-0443-2.

30. Mavroidis, M., Panagopoulou, P., Kostavasili, I., Weisleder, N., and Capetanaki, Y. (2008). A missense mutation in desmin tail domain linked to human dilated cardiomyopathy promotes cleavage of the head domain and abolishes its Z-disc localization. FASEB journal : official publication of the Federation of American Societies for Experimental Biology 22, 3318–3327. 10.1096/fj.07-088724.

31. Goldfarb, L.G., Park, K.Y., Cervenakova, L., Gorokhova, S., Lee, H.S., Vasconcelos, O., Nagle, J.W., Semino-Mora, C., Sivakumar, K., and Dalakas, M.C. (1998). Missense mutations in desmin associated with familial cardiac and skeletal myopathy. Nature genetics 19, 402–403. 10.1038/1300.

32. Tannous, P., Zhu, H., Johnstone, J.L., Shelton, J.M., Rajasekaran, N.S., Benjamin, I.J., Nguyen, L., Gerard, R.D., Levine, B., Rothermel, B.A., and Hill, J.A. (2008). Autophagy is an adaptive response in desmin-related cardiomyopathy. Proc Natl Acad Sci U S A 105, 9745–9750. 10.1073/pnas.0706802105.

33. Rawnsley, D.R., Islam, M., Zhao, C., Kargar Gaz Kooh, Y., Mendoza, A., Navid, H., Kumari, M., Guan, X., Murphy, J.T., Nigro, J., et al. (2026). Mitophagy Facilitates Cytosolic Proteostasis to Preserve Cardiac Function. Accepted at Circulation Research. 10.1101/2024.11.24.624947.

34. Pan, J.A., Sun, Y., Jiang, Y.P., Bott, A.J., Jaber, N., Dou, Z., Yang, B., Chen, J.S., Catanzaro, J.M., Du, C., et al. (2016). TRIM21 Ubiquitylates SQSTM1/p62 and Suppresses Protein Sequestration to Regulate Redox Homeostasis. Molecular cell 62, 149–151. 10.1016/j.molcel.2016.03.015.

35. Schulze, P.C., Drosatos, K., and Goldberg, I.J. (2016). Lipid Use and Misuse by the Heart. Circulation Research 118, 1736–1751. 10.1161/CIRCRESAHA.116.306842.

36. Lai, M., Amato, R., La Rocca, V., Bilgin, M., Freer, G., Spezia, P., Quaranta, P., Piomelli, D., and Pistello, M. (2021). Acid ceramidase controls apoptosis and increases autophagy in human melanoma cells treated with doxorubicin. Sci Rep 11, 11221. 10.1038/s41598-021-90219-1.

37. Xue, Q., Zhang, T., Zhu, R., Qian, Y., Dong, X., Mo, L., and Jiang, Y. (2023). Inhibition of Ceramide Synthesis Attenuates Chronic Ethanol Induced Cardiotoxicity by Restoring Lysosomal Function and Reducing Necroptosis. Alcohol Alcohol 58, 164–174. 10.1093/alcalc/agac067.

38. Chokshi, A., Drosatos, K., Cheema, F.H., Ji, R., Khawaja, T., Yu, S., Kato, T., Khan, R., Takayama, H., Knoll, R., et al. (2012). Ventricular assist device implantation corrects myocardial lipotoxicity, reverses insulin resistance, and normalizes cardiac metabolism in patients with advanced heart failure. Circulation 125, 2844–2853. 10.1161/CIRCULATIONAHA.111.060889.

39. Ji, R., Akashi, H., Drosatos, K., Liao, X., Jiang, H., Kennel, P.J., Brunjes, D.L., Castillero, E., Zhang, X., Deng, L.Y., et al. (2017). Increased de novo ceramide synthesis and accumulation in failing myocardium. JCI Insight 2. 10.1172/jci.insight.96203.

40. Tidhar, R., Zelnik, I.D., Volpert, G., Ben-Dor, S., Kelly, S., Merrill, A.H., Jr., and Futerman, A.H. (2018). Eleven residues determine the acyl chain specificity of ceramide synthases. The Journal of biological chemistry 293, 9912–9921. 10.1074/jbc.RA118.001936.

41. Mullen, T.D., Spassieva, S., Jenkins, R.W., Kitatani, K., Bielawski, J., Hannun, Y.A., and Obeid, L.M. (2011). Selective knockdown of ceramide synthases reveals complex interregulation of sphingolipid metabolism. J Lipid Res 52, 68–77. 10.1194/jlr.M009142.

42. Jaishy, B., Zhang, Q., Chung, H.S., Riehle, C., Soto, J., Jenkins, S., Abel, P., Cowart, L.A., Van Eyk, J.E., and Abel, E.D. (2015). Lipid-induced NOX2 activation inhibits autophagic flux by impairing lysosomal enzyme activity. J Lipid Res 56, 546–561. 10.1194/jlr.M055152.

43. Tong, M., Saito, T., Zhai, P., Oka, S.I., Mizushima, W., Nakamura, M., Ikeda, S., Shirakabe, A., and Sadoshima, J. (2019). Mitophagy Is Essential for Maintaining Cardiac Function During High Fat Diet-Induced Diabetic Cardiomyopathy. Circ Res 124, 1360–1371. 10.1161/CIRCRESAHA.118.314607.

44. Halbleib, K., Pesek, K., Covino, R., Hofbauer, H.F., Wunnicke, D., Hänelt, I., Hummer, G., and Ernst, R. (2017). Activation of the Unfolded Protein Response by Lipid Bilayer Stress. Molecular cell 67, 673–684.e678. 10.1016/j.molcel.2017.06.012.

45. Kurouski, D. (2023). Elucidating the Role of Lipids in the Aggregation of Amyloidogenic Proteins. Acc Chem Res 56, 2898–2906. 10.1021/acs.accounts.3c00386.

46. Pan, B., Li, J., Parajuli, N., Tian, Z., Wu, P., Lewno, M.T., Zou, J., Wang, W., Bedford, L., Mayer, R.J., et al. (2020). The Calcineurin-TFEB-p62 Pathway Mediates the Activation of Cardiac Macroautophagy by Proteasomal Malfunction. Circ Res. 10.1161/CIRCRESAHA.119.316007.

47. Zheng, Q., Su, H., Ranek, M.J., and Wang, X. (2011). Autophagy and p62 in cardiac proteinopathy. Circ.Res. 109, 296–308. CIRCRESAHA.111.244707 [pii];10.1161/CIRCRESAHA.111.244707 [doi].

48. Sergin, I., Bhattacharya, S., Emanuel, R., Esen, E., Stokes, C.J., Evans, T.D., Arif, B., Curci, J.A., and Razani, B. (2016). Inclusion bodies enriched for p62 and polyubiquitinated proteins in macrophages protect against atherosclerosis. Science signaling 9, ra2. 10.1126/scisignal.aad5614.

49. Ghosh, R., Fatahian, A.N., Rouzbehani, O.M.T., Hathaway, M.A., Mosleh, T., Vinod, V., Vowles, S., Stephens, S.L., Chung, S.D., Cao, I.D., et al. (2024). Sequestosome 1 (p62) mitigates hypoxia-induced cardiac dysfunction by stabilizing hypoxia-inducible factor 1alpha and nuclear factor erythroid 2-related factor 2. Cardiovasc Res 120, 531–547. 10.1093/cvr/cvae023.

50. Schiattarella, G.G., Altamirano, F., Tong, D., French, K.M., Villalobos, E., Kim, S.Y., Luo, X., Jiang, N., May, H.I., Wang, Z.V., et al. (2019). Nitrosative stress drives heart failure with preserved ejection fraction. Nature 568, 351–356. 10.1038/s41586-019-1100-z.

51. Tadinada, S.M., Weatherford, E.T., Collins, G.V., Bhardwaj, G., Cochran, J., Kutschke, W., Zimmerman, K., Bosko, A., O’Neill, B.T., Weiss, R.M., and Abel, E.D. (2021). Functional resilience of C57BL/6J mouse heart to dietary fat overload. American journal of physiology. Heart and circulatory physiology 321, H850–h864. 10.1152/ajpheart.00419.2021.

52. Meister-Broekema, M., Freilich, R., Jagadeesan, C., Rauch, J.N., Bengoechea, R., Motley, W.W., Kuiper, E.F.E., Minoia, M., Furtado, G.V., van Waarde, M., et al. (2018). Myopathy associated BAG3 mutations lead to protein aggregation by stalling Hsp70 networks. Nat Commun 9, 5342. 10.1038/s41467-018-07718-5.

53. Liu, L., Shi, X., Bharadwaj, K.G., Ikeda, S., Yamashita, H., Yagyu, H., Schaffer, J.E., Yu, Y.H., and Goldberg, I.J. (2009). DGAT1 expression increases heart triglyceride content but ameliorates lipotoxicity. J Biol Chem 284, 36312–36323. 10.1074/jbc.M109.049817.

54. Lemaitre, R.N., Jensen, P.N., Hoofnagle, A., McKnight, B., Fretts, A.M., King, I.B., Siscovick, D.S., Psaty, B.M., Heckbert, S.R., Mozaffarian, D., and Sotoodehnia, N. (2019). Plasma Ceramides and Sphingomyelins in Relation to Heart Failure Risk. Circulation. Heart failure 12, e005708. 10.1161/circheartfailure.118.005708.

55. Peterson, L.R., Xanthakis, V., Duncan, M.S., Gross, S., Friedrich, N., Volzke, H., Felix, S.B., Jiang, H., Sidhu, R., Nauck, M., et al. (2018). Ceramide Remodeling and Risk of Cardiovascular Events and Mortality. J Am Heart Assoc 7. 10.1161/JAHA.117.007931.

56. Mizutani, Y., Kihara, A., and Igarashi, Y. (2005). Mammalian Lass6 and its related family members regulate synthesis of specific ceramides. Biochem J 390, 263–271. 10.1042/BJ20050291.

57. Wang, X., Li, Q., Liu, Y., Jiang, H., and Chen, W. (2021). Intermittent fasting versus continuous energy-restricted diet for patients with type 2 diabetes mellitus and metabolic syndrome for glycemic control: A systematic review and meta-analysis of randomized controlled trials. Diabetes research and clinical practice 179, 109003. 10.1016/j.diabres.2021.109003.

58. Patikorn, C., Roubal, K., Veettil, S.K., Chandran, V., Pham, T., Lee, Y.Y., Giovannucci, E.L., Varady, K.A., and Chaiyakunapruk, N. (2021). Intermittent Fasting and Obesity-Related Health Outcomes: An Umbrella Review of Meta-analyses of Randomized Clinical Trials. JAMA Netw Open 4, e2139558. 10.1001/jamanetworkopen.2021.39558.

59. Sciarretta, S., Yee, D., Nagarajan, N., Bianchi, F., Saito, T., Valenti, V., Tong, M., Del Re, D.P., Vecchione, C., Schirone, L., et al. (2018). Trehalose-Induced Activation of Autophagy Improves Cardiac Remodeling After Myocardial Infarction. Journal of the American College of Cardiology 71, 1999–2010. 10.1016/j.jacc.2018.02.066.

60. Jeong, S.J., Stitham, J., Evans, T.D., Zhang, X., Rodriguez-Velez, A., Yeh, Y.S., Tao, J., Takabatake, K., Epelman, S., Lodhi, I.J., et al. (2021). Trehalose causes low-grade lysosomal stress to activate TFEB and the autophagy-lysosome biogenesis response. Autophagy, 1-13. 10.1080/15548627.2021.1896906.

61. Sergin, I., Evans, T.D., Zhang, X., Bhattacharya, S., Stokes, C.J., Song, E., Ali, S., Dehestani, B., Holloway, K.B., Micevych, P.S., et al. (2017). Exploiting macrophage autophagy-lysosomal biogenesis as a therapy for atherosclerosis. Nat Commun 8, 15750. 10.1038/ncomms15750.

62. Goldenberg, J.R., Carley, A.N., Ji, R., Zhang, X., Fasano, M., Schulze, P.C., and Lewandowski, E.D. (2019). Preservation of Acyl Coenzyme A Attenuates Pathological and Metabolic Cardiac Remodeling Through Selective Lipid Trafficking. Circulation 139, 2765–2777. 10.1161/circulationaha.119.039610.

