## Supplemental FIgures for "Impaired Proteostasis is an Early Feature of the Diabetic Heart in Humans and Mice"

**Supplementary Figures and Tables**

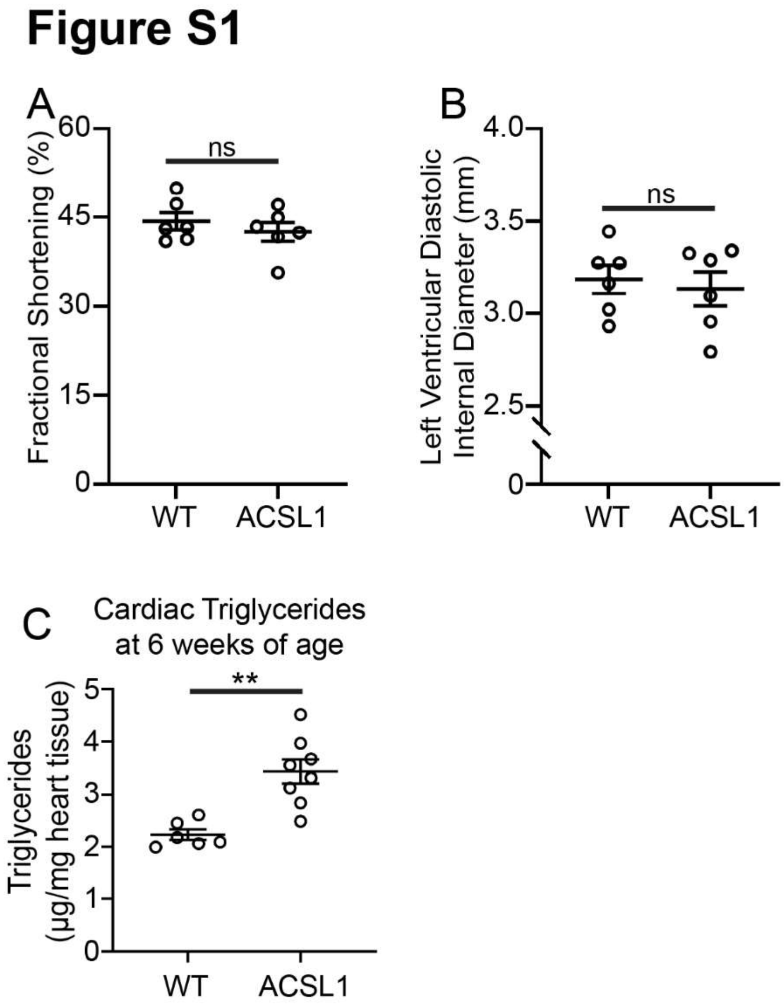


**Supplemental Figure S1. Six-week-old ACSL1 hearts exhibit normal cardiac function by echocardiography and increased triglyceride levels. (A-B)** Quantitation of left ventricular endocardial fractional shortening (A) and left ventricular diastolic internal diameter (B) in wild-type (WT) or MHC-ACSL1 transgenic mice (ACSL1) at 6 weeks of age. Statistical testing by t-test, ns $p>0.05$. **(C)** Triglyceride levels in 6-week-old wild-type (WT) and MHC-ACSL1 transgenic mouse hearts. Statistical testing by t-test, ** $p<0.01$.

### Figure S2

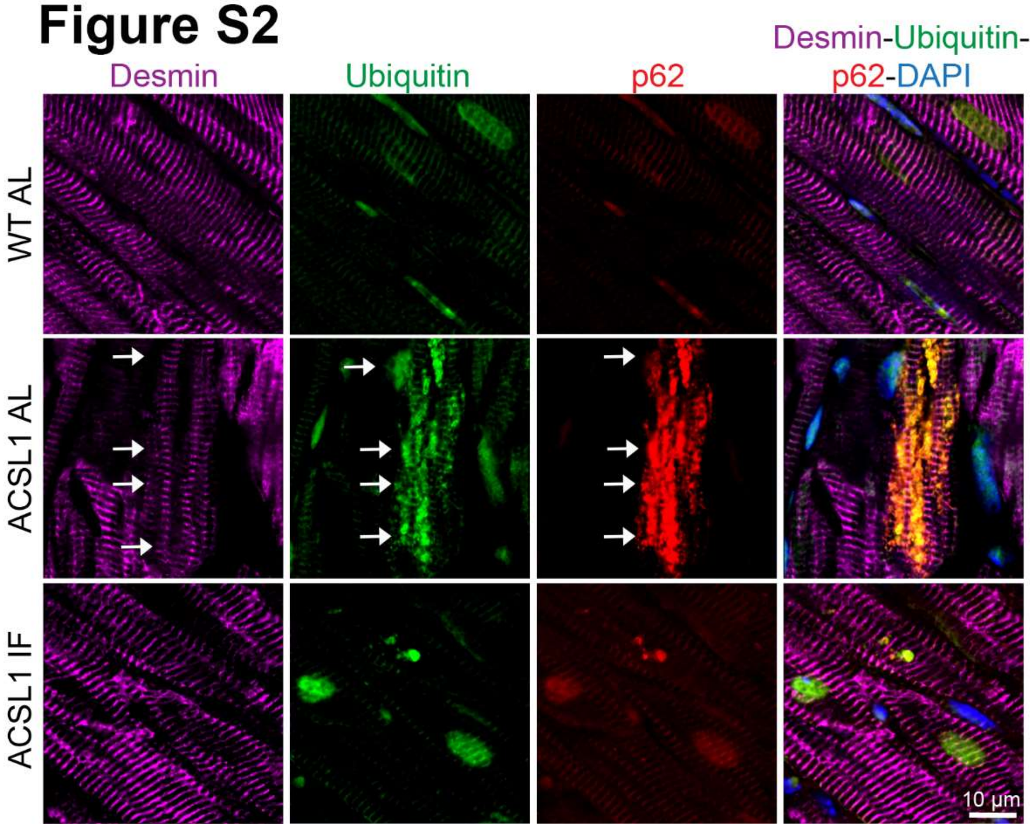


**Supplemental Figure S2. Intermittent fasting rescues protein aggregate pathology in the myocardium**. (A) Immunohistochemistry staining of 12-week-old hearts from wild-type (WT) or MHC-ACSL1 transgenic hearts under ad-libitum (AL) or intermittent fasting (IF) dietary treatment. Images show staining for Desmin (magenta), ubiquitin (green), p62 (red), and DAPI (blue). Arrows point to protein aggregates.

### Figure S3

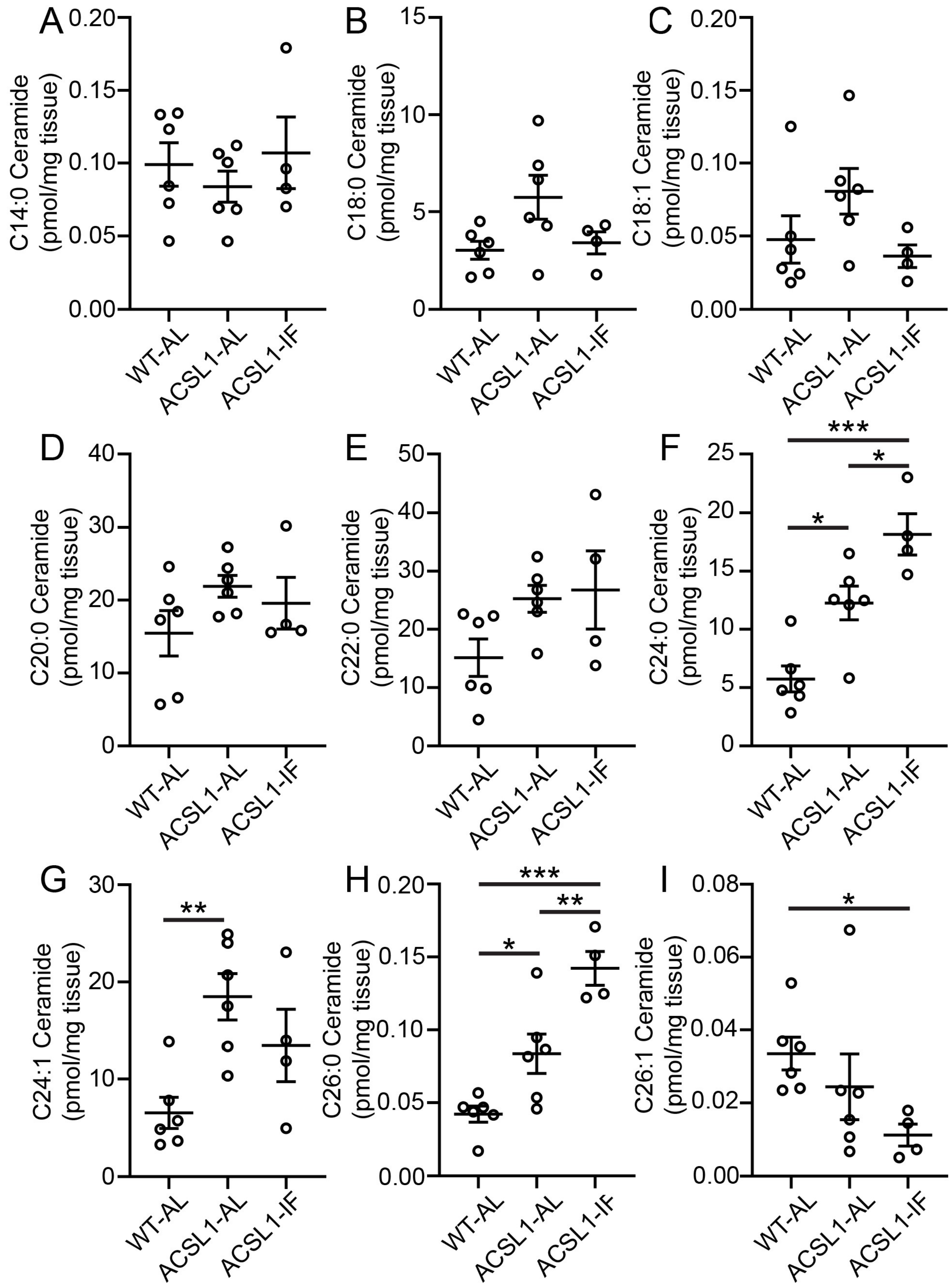
A
B
C
D
E
F
G
H
I
C14:0 Ceramide (pmol/mg tissue)
C18:0 Ceramide (pmol/mg tissue)
C18:1 Ceramide (pmol/mg tissue)
C20:0 Ceramide (pmol/mg tissue)
C22:0 Ceramide (pmol/mg tissue)
C24:0 Ceramide (pmol/mg tissue)
C24:1 Ceramide (pmol/mg tissue)
C26:0 Ceramide (pmol/mg tissue)
C26:1 Ceramide (pmol/mg tissue)
WT-AL
ACSL1-AL
ACSL1-IF
***
**
*

**Supplemental Figure S3. Effects of cardiac lipid overload in ACSL1 mouse hearts under ad-lib and intermittent fasting conditions.** (A-I) Levels of indicated ceramide species in 12-week-old mouse hearts from wild-type mice on ad-libitum diet (WT AL), ACSL1 transgenic mice on ad-libitum diet (ACSL1 AL), or ACSL1 transgenic mice after 6 weeks of every-other-day intermittent fasting (ACSL1 IF) from age 6 weeks to 12 weeks. Ceramide levels are normalized to tissue mass. Statistical testing is by one-way ANOVA followed by post-hoc testing by Tukey's test (A, B, E, F, G, H) or by Kruskal-Wallis test with post-hoc testing by Dunn's test (C, D, I). * $p < 0.05$, ** $p<0.01$, *** $P<0.001$.

### Figure S4

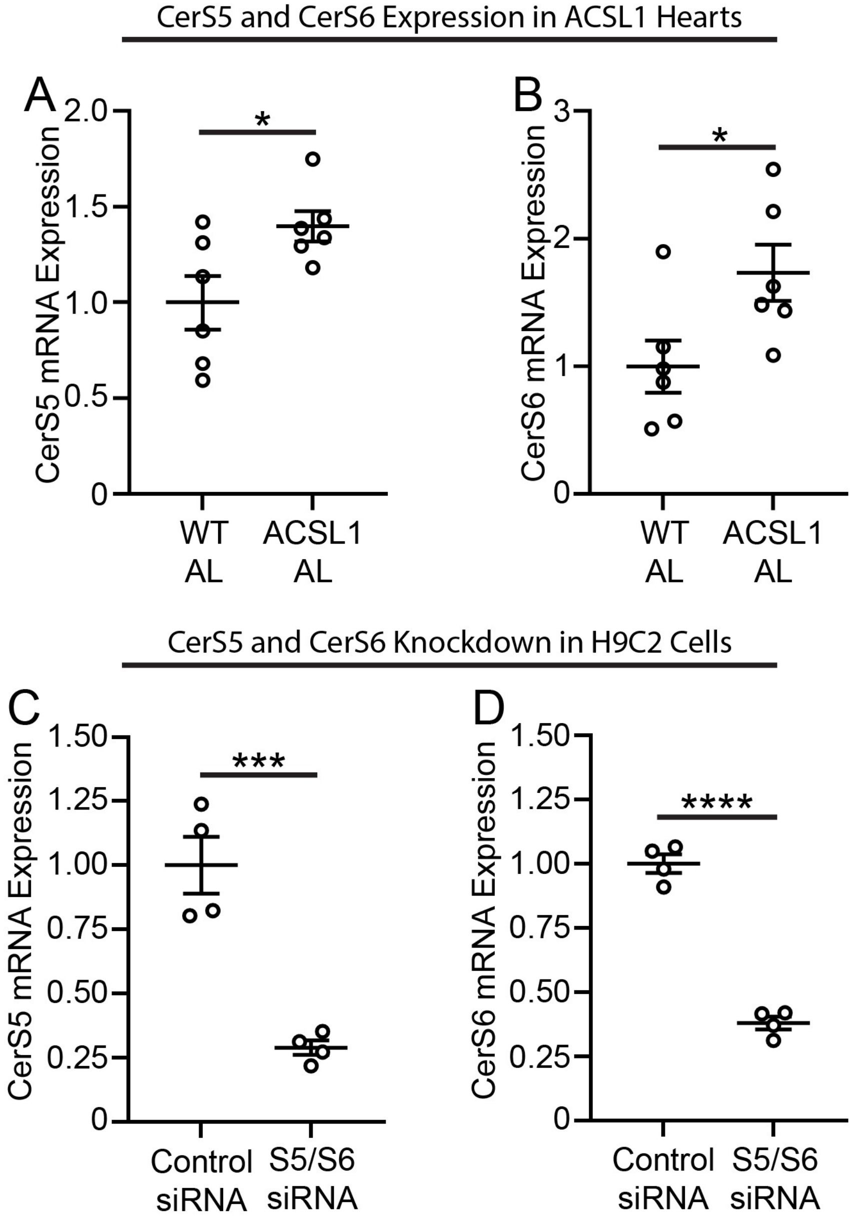


**Supplemental Figure S4. Ceramide synthase 5 and 6 mRNA expression in the hearts of MHC-ACSL1 transgenic mice, and ceramide synthase expression in H9C2 cells following siRNA targeting. (A-B)** Expression of ceramide synthase 5 (CerS5) and ceramide synthase 6 (CerS6) as assessed by quantitative RT-PCR in 6-week-old wild type and MHC-ACSL1 hearts. All mice are on ad-libitum diets (AL). **(C-D)** CerS5 and CerS6 mRNA levels in rat H9C2 cells 48 hours after treatment with siRNA against rat CerS5 and CerS6 (S5/S6), as compared to cells treated with non-targeting control siRNA. Statistical comparisons in A-D are by unpaired t-test. * $p<0.05$, *** $p<0.001$, **** $p<0.0001$.

### Figure S5

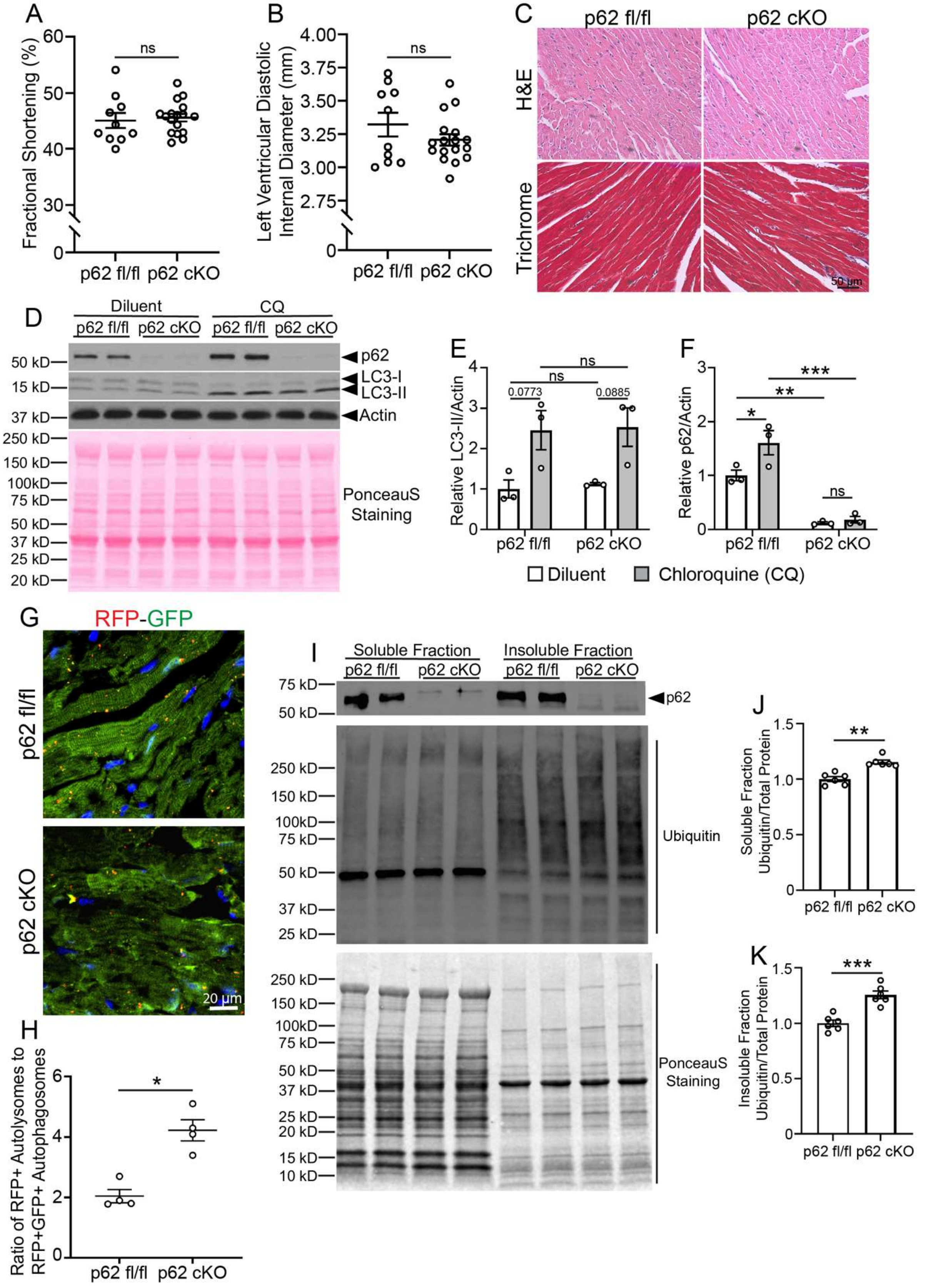
A
Fractional Shortening (%)
ns
p62 fl/fl
p62 cKO
B
Left Ventricular Diastolic Internal Diameter (mm)
ns
p62 fl/fl
p62 cKO
C
p62 fl/fl
p62 cKO
H&E
Trichrome
50 µm
D
Diluent
CQ
p62 fl/fl
p62 cKO
p62 fl/fl
p62 cKO
p62
LC3-I
LC3-II
Actin
PonceauS Staining
E
Relative LC3-II/Actin
0.0773
0.0885
ns
ns
p62 fl/fl
p62 cKO
F
Relative p62/Actin
*
**
***
ns
p62 fl/fl
p62 cKO
Diluent
Chloroquine (CQ)
G
RFP-GFP
p62 fl/fl
p62 cKO
20 µm
H
Ratio of RFP+ Autolysomes to RFP+GFP+ Autophagosomes
*
p62 fl/fl
p62 cKO
I
Soluble Fraction
Insoluble Fraction
p62 fl/fl
p62 cKO
p62 fl/fl
p62 cKO
p62
Ubiquitin
PonceauS Staining
J
Soluble Fraction Ubiquitin/Total Protein
**
p62 fl/fl
p62 cKO
K
Insoluble Fraction Ubiquitin/Total Protein
***
p62 fl/fl
p62 cKO

**Supplementary Figure S5. Cardiac myocyte specific p62 ablation does not impair cardiac structure, function, or autophagic flux in young mice. (A-B)** %Left ventricular endocardial fractional shortening (A) and left ventricular diastolic internal diameter (B) in 8-week-old Myh6-Cre p62 flox/flox (p62 cKO) or in p62 flox/flox (p62 fl/fl) control mice. Statistical testing by t-test. **(C)** Hematoxylin and eosin (H&E) and Masson's trichrome staining in 12-week-old p62 fl/fl and p62 cKO hearts. **(D)** Immunoblotting in 8-week-old p62 fl/fl and p62 cKO mice treated with IP chloroquine (CQ) injection (dose-80 mg/kg) or diluent control injection 4 hours prior to sacrifice. **(E-F)** Quantitation of LC3B-II (E) and p62 (F) from panel D, normalized to actin expression. Statistical testing by two-way ANOVA with Tukey's test for multiple comparison testing. **(G)** Fluorescent imaging of RFP and GFP expression from frozen sections from p62 fl/fl and p62 cKO mice carrying the CAG-LC3-RFP-GFP transgenic reporter allele. **(H)** Quantitation of ratio of RFP+ autolysosomes to RFP+ GFP+ autophagosomes per high power field from panel G. Puncta were quantified from five separate images per sample and then averaged. Statistical comparison by t-test. **(I)** Immunoblotting of the NP-40 soluble and NP-40 insoluble fractions from 8-week-old p62 fl/fl or p62 cKO hearts. **(J-K)** Quantitation of ubiquitin levels in the soluble (J) and insoluble (K) fractions from panel I. Ubiquitin levels are normalized to Ponceau S staining. Statistical comparison by t-test. For statistical comparisons in this figure, ns $p>0.05$, ** $p<0.01$, *** $p<0.001$.

### Figure S6

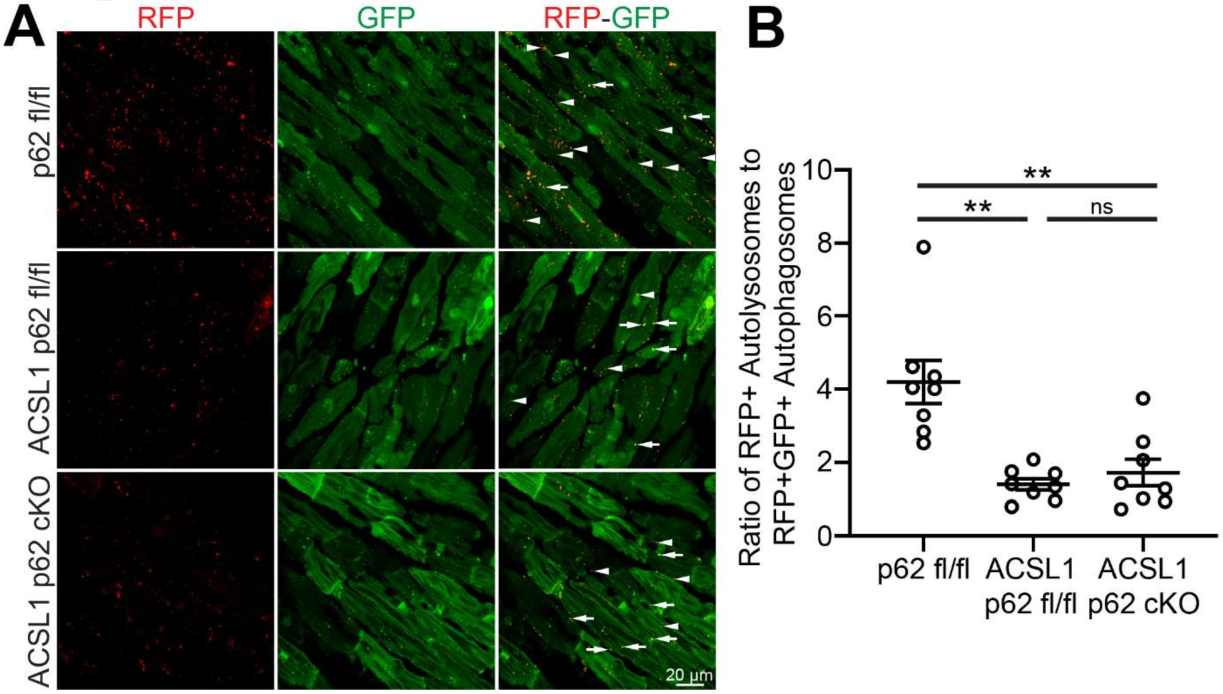


**Supplemental Figure S6. Deletion of p62 in cardiomyocytes does not alter the impairment of autophagic flux in cardiomyocytes in ACSL1 transgenic mice. (A)** Fluorescent imaging of RFP and GFP expression from frozen sections from 10-week-old hearts from p62 fl/fl vs. MHC-ACSL1 p62 fl/fl vs. MHC-ACSL1 p62 cKO mice. All mice were heterozygous for the CAG-LC3-RFP-GFP transgenic reporter allele. White arrowheads indicate RFP+ GFP – autolysosomes; white arrows indicate double RFP+ GFP+ autophagosomes. **(B)** Ratio of RFP+ GFP- puncta (autolysosomes) to RFP+GFP+ puncta (autophagosomes) from Panel A. Puncta were scored from five images per sample and then averaged. Statistic comparison by one-way ANOVA with Tukey's test for multiple comparison testing. For statistical comparisons in this figure, ns $p>0.05$, ** $p<0.01$.

### Figure S7

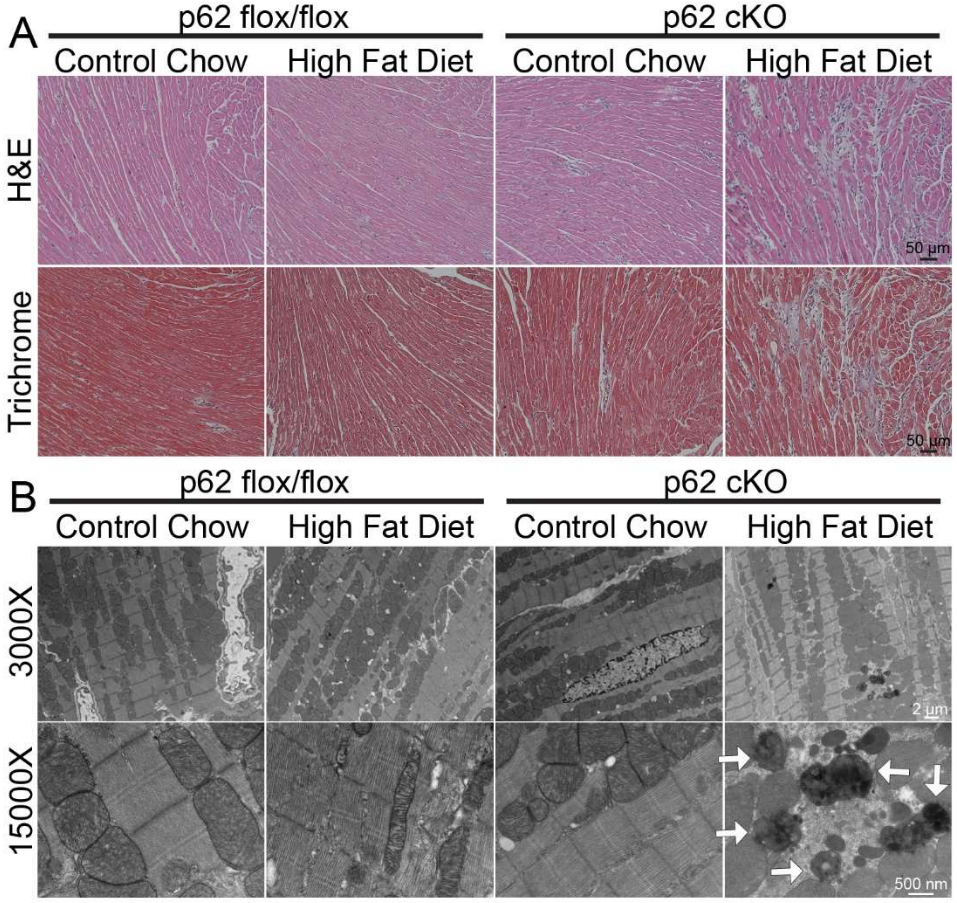


**Supplementary Figure S7. Cardiac myocyte ablation of p62 induces fibrosis and protein aggregates in mice fed a high-fat diet. (A)** Representative hematoxylin and eosin-stained myocardial sections from 32 week-old p62 fl/fl or p62 cKO mice fed 24 weeks of control chow ad-lib (Chow) or a high-fat diet ad-lib diet (HFD). **(B)** Representative transmission electron micrographs depicting myocardial ultrastructure from mice treated as in A. Arrows point to large electron dense protein aggregates.

**Supplemental Table S1: Morphometric data for WT or MHC-ACSL1 mice on indicated diets at 12 weeks of age.**

| | **Wild-Type Mice Ad-lib Diet (n=24)** | **MHC-ACSL1 Mice Ad-lib Diet (n=25)** | **MHC-ACSL1 Mice IF Diet (n=17)** |
|---|---|---|---|
| **BW (g)** | 21.50 ± 0.36 | 21.77 ± 0.45 | 20.58 ± 0.46 |
| **HW/BW (mg/g)** | 4.85 ± 0.05 | 6.66 ± 0.23 **** | 5.30 ± 0.09 * $^{###}$ |
| **HW/TL (g/mm)** | 6.26 ± 0.13 | 8.77 ± 0.32 **** | 6.66 ± 0.14 $^{####}$ |

Data represent mean ± SEM. Statistical testing by one-way ANOVA followed by post-hoc Tukey's test (for BW) or by non-parametric Kruskal-Wallis test followed by post-hoc testing with Dunn's test (for HW/BW, HW/TL). * $p<0.05$, **** $p<0.0001$ for comparison vs. Wild-type Ad-lib Diet group; $^{###}$ $p<0.001$, $^{####}$ $p<0.0001$ for comparison vs. MHC-ACSL1 Ad-lib diet group. All other statistical comparisons did not meet significance ($p>0.05$). BW = Body Weight, HW = Heart Weight, TL = Tibia Length.

**Supplemental Table S2: Echocardiographic data for WT or MHC-ACSL1 mice on indicated diets at 12 weeks of age.**

| | **Wild-Type Mice Ad-lib Diet (n=4)** | **MHC-ACSL1 Mice Ad-lib Diet (n=9)** | **MHC-ACSL1 Mice IF Diet (n=11)** |
|---|---|---|---|
| **LVIDd (mm)** | 3.39 ± 0.08 | 4.19 ± 0.14 * | 3.45 ± 0.06 $^{\#\#}$ |
| **LVIDs (mm)** | 1.95 ± 0.05 | 3.39 ± 0.18 *** | 2.31 ± 0.10 $^{\#\#\#}$ |
| **FS (%)** | 42.33 ± 0.86 | 19.26 ± 1.86 **** | 33.86 ± 1.69 $^{\#\#\#}$ |
| **LV mass (g)** | 130.96 ± 6.84 | 147.62 ± 4.30 | 113.10 ± 5.66 $^{\#\#}$ |
| **HR (beats per min)** | 646.4 ± 14.1 | 553.9 ± 7.2 *** | 546.4 ± 5.8 **** |

Data represent mean ± SEM. Statistical testing by one-way ANOVA followed by post-hoc Tukey's test (for LVIDs, FS, LV Mass, HR) or by non-parametric Kruskal-Wallis test followed by post-hoc testing with Dunn's test (for LVIDd). * p<0.05, *** p<0.001, **** p<0.0001 for comparison vs. Wild-Type Ad-lib Diet group; $^{\#\#}$ p<0.01, $^{\#\#\#}$ p<0.001 for comparison vs. MHC-ACSL1 Ad-lib diet group. All other statistical comparisons did not meet significance (p>0.05). LVIDd = Left Ventricular Internal Diameter End Diastole, LVIDs = Left Ventricular Internal Diameter End Systole, FS = Fractional Shortening, LV mass = Left-Ventricular mass based on M-mode echocardiography measurements, HR = Heart rate.

**Supplemental Table S3: Morphometric data for mice of indicated genotypes at 6 weeks of age.**

| | **p62 fl/fl mice (n=10)** | **MHC-ACSL1 p62 fl/fl mice (n=10)** | **MHC-ACSL1 p62 cKO mice (n=8)** |
|---|---|---|---|
| **BW (g)** | 17.75 ± 0.39 | 17.88 ± 0.41 | 18.38 ± 0.58 |
| **HW/BW (mg/g)** | 4.79 ± 0.10 | 6.18 ± 0.10 **** | 6.27 ± 0.09 *** |
| **HW/TL (g/mm)** | 5.60 ± 0.19 | 7.25 ± 0.19 **** | 7.54 ± 0.22 **** |

Data represent mean ± SEM. Statistical testing by one-way ANOVA followed by post-hoc Tukey's test (for BW, HW/TL) or by non-parametric Kruskal-Wallis test followed by post-hoc testing with Dunn's test (for HW/BW). *** $p<0.001$, **** $p<0.0001$ for comparison vs. p62 fl/fl group; there are no statistically significant differences between MHC-ACSL1 p62 fl/fl and MHC-ACSL1 p62 cKO groups. All other statistical comparisons did not meet significance ($p>0.05$). BW = Body Weight, HW = Heart Weight, TL = Tibia Length.

**Supplemental Table S4: Echocardiographic data for mice of indicated genotypes at 6 weeks of age.**

| | **p62 fl/fl mice (n=8)** | **MHC-ACSL1 p62 fl/fl mice (n=8)** | **MHC-ACSL1 p62 cKO mice (n=8)** |
|---|---|---|---|
| **LVIDd (mm)** | 3.21 ± 0.08 | 3.45 ± 0.05 * | 3.41 ± 0.04 |
| **LVIDs (mm)** | 1.86 ± 0.07 | 2.27 ± 0.5 *** | 2.26 ± 0.04 *** |
| **FS (%)** | 41.96 ± 1.40 | 34.35 ± 0.66 *** | 33.93 ± 0.90 *** |
| **LV mass (g)** | 68.71 ± 2.24 | 85.70 ± 3.03 *** | 81.37 ± 1.34 ** |
| **HR (beats per min)** | 587.9 ± 14.6 | 547.6 ± 5.9 * | 551.7 ± 7.3 |

Data represent mean ± SEM. Statistical testing by one-way ANOVA followed by post-hoc Tukey's test. * $p<0.05$, ** $p<0.01$, *** $p<0.001$ for comparison vs. p62 fl/fl mice group. All other statistical comparisons did not meet significance ($p>0.05$); there are no statistically significant differences between MHC-ACSL1 p62 fl/fl and MHC-ACSL1 p62 cKO groups. LVIDd = Left Ventricular Internal Diameter End Diastole, LVIDs = Left Ventricular Internal Diameter End Systole, FS = Fractional Shortening, LV mass = Left-Ventricular mass based on M-mode echocardiography measurements, HR = Heart rate.

**Supplemental Table S5: Morphometric data for mice of indicated genotypes at 10 weeks of age.**

| | **MHC-ACSL1 p62 fl/fl mice (n=11)** | **MHC-ACSL1 p62 cKO mice (n=11)** |
|---|---|---|
| **BW (g)** | 19.48 ± 0.69 | 20.51 ± 0.67 |
| **HW/BW (mg/g)** | 6.48 ± 0.17 | 6.16 ± 0.12 |
| **HW/TL (g/mm)** | 8.43 ± 0.32 | 7.91 ± 0.26 |

Data represent mean ± SEM. Statistical testing by unpaired t-test. All statistical comparisons did not meet significance ($p>0.05$). BW = Body Weight, HW = Heart Weight, TL = Tibia Length.

**Supplemental Table S6: Echocardiographic data for mice of indicated genotypes 10 weeks of age.**

| | **MHC-ACSL1 p62 fl/fl mice (n=4)** | **MHC-ACSL1 p62 cKO mice (n=6)** |
|---|---|---|
| **LVIDd (mm)** | 3.68 ± 0.17 | 4.78 ± 0.11 * |
| **LVIDs (mm)** | 2.81 ± 0.14 | 3.86 ± 0.17 ** |
| **FS (%)** | 23.53 ± 0.14 | 12.83 ± 1.93 * |
| **LV mass (g)** | 104.20 ± 6.72 | 125.30 ± 5.3 * |
| **HR (beats per min)** | 556.2 ± 4.3 | 536.9 ± 25.37 |

Data represent mean ± SEM. Statistical testing by unpaired t-test. * $p<0.05$, ** $p<0.01$ for comparison vs. MHC-ACSL1 p62 fl/fl group. All other statistical comparisons did not meet significance ($p>0.05$). LVIDd = Left Ventricular Internal Diameter End Diastole, LVIDs = Left Ventricular Internal Diameter End Systole, FS = Fractional Shortening, LV mass = Left-Ventricular mass based on M-mode echocardiography measurements, HR = Heart rate.

**Supplemental Table S7: Echocardiographic data for MHC-ACSL1 p62 cKO mice on indicated diets at 10 weeks of age.**

| | **MHC-ACSL1 p62 cKO Ad Lib Diet (n=7)** | **MHC-ACSL1 p62 cKO IF Diet (n=7)** |
|---|---|---|
| **LVIDd (mm)** | 3.96 ± 0.10 | 3.41 ± 0.10 *** |
| **LVIDs (mm)** | 3.32 ± 0.13 | 2.29 ± 0.08 **** |
| **FS (%)** | 16.39 ± 1.19 | 37.47 ± 1.06 **** |
| **LV mass (g)** | 94.20 ± 4.43 | 67.84 ± 3.07 *** |
| **HR (beats per min)** | 541.6 ± 4.2 | 585.3 ± 9.5 *** |

Data represent mean ± SEM. Statistical testing by unpaired t-test (for LVIDs, FS) and by Mann-Whitney test (for LVIDd, LV Mass, HR). *** $p<0.001$, **** $p<0.0001$ for comparison vs. MHC-ACSL1 cKO Ad Lib diet group. LVIDd = Left Ventricular Internal Diameter End Diastole, LVIDs = Left Ventricular Internal Diameter End Systole, FS = Fractional Shortening, LV mass = Left-Ventricular mass based on M-mode echocardiography measurements, HR = Heart rate.

**Supplemental Table S8: Echocardiographic data for MHC-ACSL1 p62 fl/fl vs MHC-ACSL1 p62 cKO mice on intermittent fasting at 28 weeks of age.**

| | **MHC-ACSL1 p62 fl/fl IF Diet (n=8)** | **MHC-ACSL1 p62 cKO IF Diet (n=6)** |
|---|---|---|
| **LVIDd (mm)** | 3.66 ± 0.06 | 4.35 ± 0.17 *** |
| **LVIDs (mm)** | 2.58 ± 0.08 | 3.71 ± 0.22 *** |
| **FS (%)** | 29.63 ± 1.29 | 15.11 ± 1.78 **** |
| **LV mass (g)** | 89.80 ± 4.65 | 105.17 ± 2.94 |
| **HR (beats per min)** | 566.0 ± 11.7 | 550.5 ± 13.7 |

Data represent mean ± SEM. Statistical testing by unpaired t-test (LVIDd, LVIDs, FS, LV Mass) and by Mann-Whitney test (HR). *** $p<0.001$, **** $p<0.0001$ for comparison vs. MHC-ACSL1 p62 fl/fl IF diet group. All other statistical comparisons did not meet significance ($p>0.05$). LVIDd = Left Ventricular Internal Diameter End Diastole, LVIDs = Left Ventricular Internal Diameter End Systole, FS = Fractional Shortening, LV mass = Left-Ventricular mass based on M-mode echocardiography measurements, HR = Heart rate.

**Supplemental Table S9: Morphometric data for p62 fl/fl or p62 cKO mice at 32 weeks of age following 24 weeks of high-fat diet or chow control diet.**

| | **p62 fl/fl**<br><br>**Chow Diet (n=7)** | **p62 fl/fl**<br><br>**HFD (n=7)** | **p62 cKO**<br><br>**Chow Diet (n=7)** | **p62 cKO**<br><br>**HFD (n=5)** |
|---|---|---|---|---|
| **BW (g)** | 25.40 ± 0.75 | 39.08 ± 2.60** | 25.9 ± 1.05 | 38.12 ± 3.56## |
| **BW/TL (g/mm)** | 1.48 ± 0.04 | 2.26 ± 0.15** | 1.52 ± 0.07 | 2.24 ± 0.22## |
| **HW/TL (mg/mm)** | 6.69 ± 0.19 | 7.78 ± 0.42* | 6.63 ± 0.21 | 8.11 ± 0.37# |
| **LW/TL (mg/mm)** | 8.80 ± 0.31 | 10.14 ± 1.11 | 8.37 ± 0.23 | 11.09 ± 1.42 |

Data represent mean ± SEM. Statistical testing by two-way ANOVA followed by post-hoc Tukey's test. * $p<0.05$, ** $p<0.01$ for comparison vs. p62 fl/fl Chow diet group; # $p<0.05$, ## $p<0.01$ vs. p62 cKO Chow diet group. All other statistical comparisons did not meet significance ($p>0.05$). BW = Body Weight, HW = Heart Weight, LW = Lung weight, TL = Tibia Length, HFD = High-fat diet.

**Supplemental Table S10: Echocardiographic data for p62 fl/fl or p62 cKO mice at 32 weeks of age following 24 weeks of high-fat diet or chow control diet.**

| | **p62 fl/fl Chow Diet (n=7)** | **p62 fl/fl HFD (n=7)** | **p62 cKO Chow Diet (n=7)** | **p62 cKO HFD (n=6)** |
|---|---|---|---|---|
| **LVIDd (mm)** | 3.39 ± 0.10 | 3.49 ± 0.13 | 3.53 ± 0.08 | 4.02 ± 0.28 |
| **LVIDs (mm)** | 1.97 ± 0.11 | 1.91 ± 0.08 | 2.18 ± 0.07 | 3.03 ± 0.40 ** # ‡‡ |
| **FS (%)** | 42.23 ± 1.84 | 45.14 ± 1.18 | 38.27 ± 0.79 | 26.2 ± 4.02 *** ###### ‡‡ |
| **LV mass (g)** | 84.57 ± 4.61 | 96.114 ± 7.0 | 86.90 ± 3.50 | 100.71 ± 4.98 * |
| **HR (beats per min)** | 591 ± 15.5 | 621 ± 14.2 | 650 ± 10.0 | 604 ± 26.6 |

Data represent mean ± SEM. Statistical testing by two-way ANOVA followed by post-hoc Tukey's test. * p<0.05, ** p<0.01, *** p<0.001 for comparison vs. p62 fl/fl Chow diet group; # p<0.05, #### p<0.0001 for comparison vs. p62 fl/fl HFD group; ‡‡ p<0.01 for comparison vs. p62 cKO Chow diet group. All other statistical comparisons did not meet significance (p>0.05). LVIDd = Left Ventricular Internal Diameter End Diastole, LVIDs = Left Ventricular Internal Diameter End Systole, FS = Fractional Shortening, LV mass = Left-Ventricular mass based on M-mode echocardiography measurements, HR = Heart rate.

**Supplementary Table S11: Characteristics of individuals for data reported in biochemical analysis of human ventricular myocardium (in Figure 8A-C).**

| | **Non-Diabetic Control (n=3)** | **Diabetic (n=3)** | **P-value** |
|---|---|---|---|
| **Age** | 52.0 ± 2.8 | 66.7 ± 3.3 | 0.051 |
| **Diabetes (n)** | 0 | 3 | N/A |
| **Insulin Use (n)** | 0 | 3 | N/A |
| **LV Mass (g)** | 239.7 ± 20.9 | 396.5 ± 30.9 | 0.0566 |
| **Heart Weight (g)** | 387.7 ± 25.5 | 683.5 ± 49.9 | 0.0301 |
| **Body Weight (kg)** | 107 ± 17.2 | 97.0 ± 4.0 | 0.669 |
| **BSA (m2)** | 2.27 ± 0.17 | 2.15 ± 0.13 | 0.651 |
| **LVMI (g/m2)** | 105.4 ± 1.5 | 176.5 ± 11.0 | 0.016 |
| **HMI (g/m2)** | 171.3 ± 2.9 | 304.2 ± 17.48 | 0.010 |
| **LVEF (%)** | 62.5 ± 1.44 | 63.3 ± 1.36 | 0.79 |

Data represent mean ± SEM. Statistical testing is by t-test.

**Supplementary Table S12: Characteristics of individuals for data reported in histologic analysis of human ventricular myocardium (Figure 8D-F).**

| | **Non-Diabetic Control (n=3)** | **Diabetic (n=3)** | **P-value** |
|---|---|---|---|
| **Age** | 66.0 ± 3.1 | 59.3 ±7.3 | 0.954 |
| **Diabetes (n)** | 0 | 3 | N/A |
| **Body Weight (kg)** | 73.3 ± 5.4 | 93.3 ± 6.8 | 0.835 |
| **BMI** | 24.8 ± 1.9 | 28.9 ± 0.6 | 0.995 |
| **LVEF (%)** | 67.8 ± 2.23 | 65.0 ± 2.36 | 0.79 |

Data represent mean ± SEM. Statistical testing is by t-test.
